# The *Drosophila* anterior-posterior axis is polarized by asymmetric myosin activation

**DOI:** 10.1101/2021.04.21.440791

**Authors:** Hélène Doerflinger, Vitaly Zimyanin, Daniel St Johnston

## Abstract

The *Drosophila* anterior-posterior (AP) axis is specified at mid-oogenesis when Par-1 kinase is recruited to the posterior cortex of the oocyte, where it polarises the microtubule cytoskeleton to define where the axis determinants, *bicoid* and *oskar* mRNAs localise. This polarity is established in response to an unknown signal from the follicle cells, but how this occurs is unclear. Here we show that the myosin chaperone, Unc-45 and Non-Muscle Myosin II (MyoII) are required in the germ line upstream of Par-1 in polarity establishment. Furthermore, the Myosin regulatory Light Chain (MRLC) is di-phosphorylated at the oocyte posterior in response to the follicle cell signal, inducing longer pulses of myosin contractility at the posterior and increased cortical tension. Over-expression of MRLC-T21A that cannot be di-phosphorylated or acute treatment with the Myosin light chain kinase inhibitor ML-7 abolish Par-1 localisation, indicating that posterior of MRLC di-phosphorylation is essential for polarity. Thus, asymmetric myosin activation polarizes the anterior-posterior axis by recruiting and maintaining Par-1 at the posterior cortex. This raises an intriguing parallel with AP axis formation in *C. elegans* where MyoII is also required to establish polarity, but functions to localize the anterior PAR proteins rather than PAR-1.

## Introduction

In many organisms, the primary body axis is defined by the polarisation of the egg or zygote, generating cellular asymmetries that lead to the localisation and segregation of cytoplasmic determinants. This has been extensively characterised in *C. elegans*, where the posterior pole is defined by the site of sperm entry into the unfertilised egg [1]. Polarity establishment starts when Aurora A associated with the sperm centrosome inhibits myosin activity at the posterior cortex to trigger a contraction of cortical actomyosin towards the anterior [2–4].

The anterior polarity proteins PAR-3, PAR-6 and aPKC, which are initially localised all around the egg membrane, are carried to the anterior by this cortical flow, allowing the posterior polarity factors, PAR-2, PAR-1 and Lgl to localise to the posterior cortex [5–7]. After this establishment phase, polarity is maintained by mutual antagonism between the anterior and posterior PAR proteins. The localised PAR proteins control spindle orientation and the asymmetric localisation of determinants to drive an asymmetric first cell division to produce a large anterior AB cell and a smaller posterior P cell [8].

Like *C. elegans*, *Drosophila* sets up its anterior-posterior axis at the one cell stage, but in this case, during the development of the oocyte [9]. Anterior-posterior asymmetry arises in the germarium when the oocyte moves to the posterior end of the sixteen cell germline cyst as a result of differential adhesion between the oocyte and the somatic follicle cells [10–12]. The follicle cells at the two ends of the egg chamber subsequently become specified as terminal follicle cells, rather than main body follicle cells as a result of Unpaired signalling from a pair of polar cells at each pole of the egg chamber [13, 14]. At stage 6 of oogenesis, the EGF-like ligand, Gurken, is secreted from the posterior of the oocyte to induce the adjacent terminal follicle cells at this end of the egg chamber to adopt a posterior fate instead of the default anterior fate, and these cells then signal back to the oocyte to induce its polarisation along the future anterior-posterior axis [15, 16]. Despite extensive searches, however, the polarising signal from the follicle cells has not been identified [17].

The first sign of the anterior-posterior polarisation of the *Drosophila* oocyte is the recruitment of the Par-1 kinase to the posterior cortex at stage 7 of oogenesis, in a process that depends on the actin cytoskeleton [18–20]. At the same time, aPKC and Par-6 are excluded from the posterior cortex, while the Par-3 orthologue, Baz, disappears from the posterior slightly later. This cortical polarity is then maintained by mutual antagonism between the anterior and posterior Par proteins, in which Par-1 phosphorylates Baz to exclude it from the posterior cortex and aPKC phosphorylates Par-1 to prevent it from localising laterally [21, 22]. Par-1 transduces this cortical polarity to the microtubule cytoskeleton by repressing the formation of noncentrosomal microtubule organising centres (ncMTOCs) posteriorly, leading to the formation of a weakly-polarised microtubule network that directs the kinesin-dependent transport of the posterior determinant, *oskar* mRNA, to the posterior pole [23, 24]}. Almost nothing is known about how this Par protein asymmetry is established, except that the Ubiquitin ligase, Slimb, is necessary for the posterior recruitment of Par-1 [25]. Here we report that polarity signalling induces the specific activation of non-muscle Myosin II (MyoII) at the posterior of the oocyte and show that this acts upstream of Slimb in the recruitment of Par-1, making it the first sign of polarity establishment identified to date.

## Results

We identified a complementation group of three alleles that we named *poulpe* in a germline clone screen for mutants that disrupt the posterior localisation of GFP-Staufen, which acts as a marker for *oskar* mRNA [26]. In wild-type stage 9-10 egg chambers, Staufen and *oskar* mRNA localise to a well-defined crescent at the posterior of the oocyte, whereas they are often not localised at all, or localise to the centre of the oocyte in *poulpe* homozygous mutant germline clones (Figure 1A-C, E-G and Figure S1). In some weaker cases, Staufen and *oskar* mRNA reach the posterior region, but form a diffuse cloud rather than a crescent, which is reminiscent of phenotype seen in mutants that fail to anchor Staufen/*oskar* mRNA complexes once they are localised (Figure 1D and H and Figure S1).

**Figure 1.**
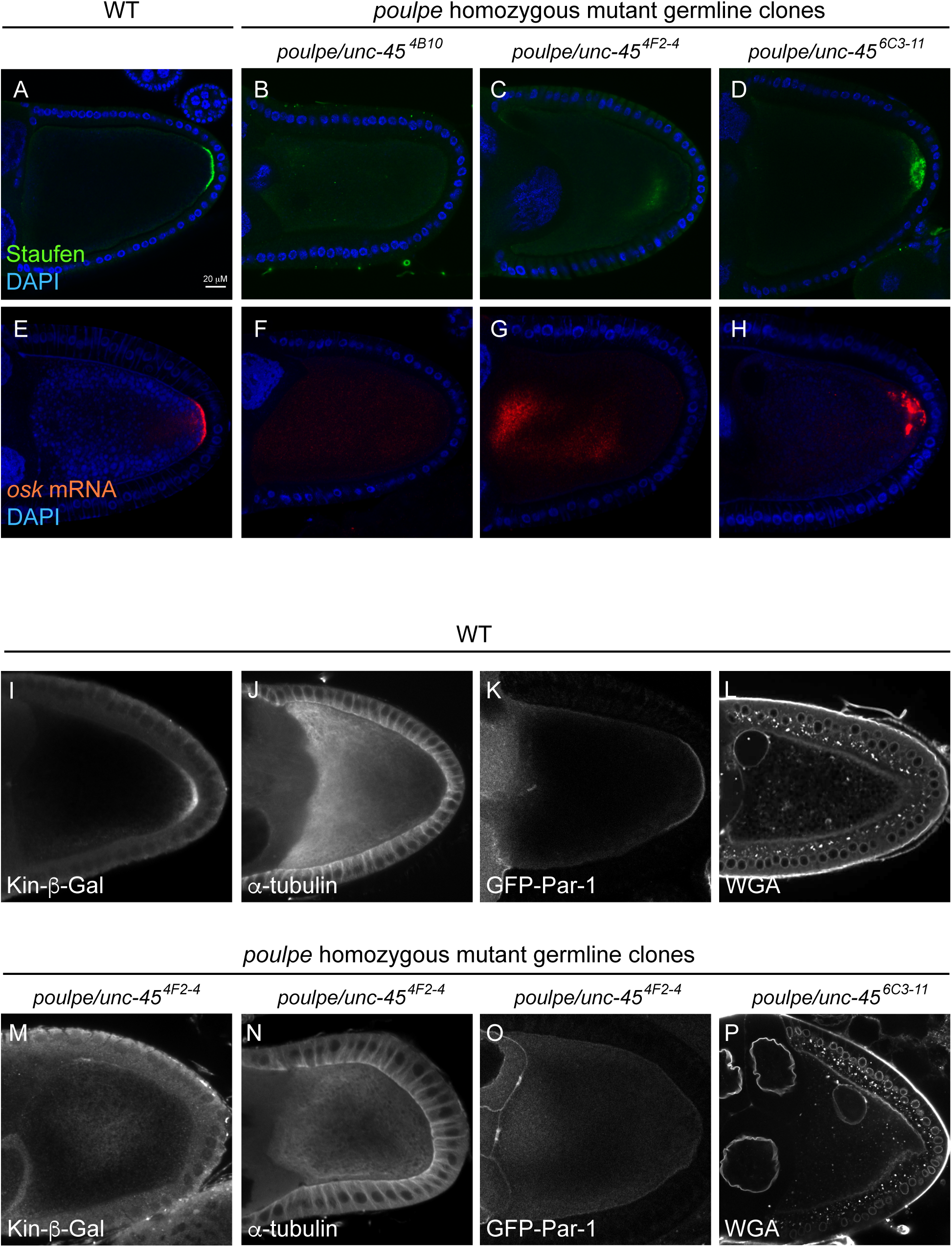
The *poulpe* gene is required for the oocyte polarisation and acts uptreams of Par-1. (A) A confocal image of a wild-type egg chamber showing the localisation of Staufen (green) in a crescent at the posterior cortex of the oocyte; DAPI (blue). (B-D) Confocal images showing examples of Staufen localisation in *poulpe* homozygous mutant germline clones. Staufen is either diffusely localised (D), localised to the centre of the oocyte (C) or forms a diffuse cloud near the posterior pole; Staufen (green) and DAPI (blue). (E) A confocal image of a wild-type egg chamber showing the localisation of *oskar* mRNA (red) to the posterior cortex of the oocyte; DAPI (blue). (F and H) Confocal images showing examples of *oskar* mRNA localisation in *poulpe* homozygous mutant germline clones; *oskar* mRNA (red) and DAPI (blue). (I and J) Kinesin-β−galactosidase localisation in wild-type (I) and *poulpe*^4F2-4^ mutant (J) oocytes. The constitutively-active Kinesin-β−galactosidase (white) localises to the posterior cortex of the wildtype oocyte by moving towards microtubule plus ends. Kinesin-β−galactosidase localises to the centre of the *poulpe*^42F4^ mutant oocyte, indicating that the oocyte is not polarised and the plus ends are not focussed on the posterior. (K and L) *α*-tubulin staining showing the microtubule organisation in wild-type (K) and *poulpe*^4F2-4^ mutant (L) oocytes. In the wild-type oocyte, the microtubules form an anterior to posterior gradient and are most dense along the anterior and lateral cortex where their stable minus ends are anchored. In the *poulpe*^4F2-4^ mutant, the microtubules are nucleated all around the oocyte cortex, forming a density gradient from the cortex to the centre. (M and N) Confocal images showing the localisation of Par-1-GFP expressed from a protein trap line in wild-type (M) and *poulpe*^4F2-4^ mutant (N) oocytes. Par-1 forms a crescent at the posterior of the wild-type oocyte, but is not localised in the *poulpe*^4F2-4^ mutant. (O and P) Confocal images showing Wheat germ agglutinin (WGA) staining to label the nuclei in a wild-type (O) and *poulpe*^6C3-11^ egg chamber. The oocyte nucleus is anchored at the dorsal/anterior corner of the wild-type oocyte, but is mislocalised to the lateral cortex in the *poulpe*^6C3-11^ mutant.

Since the strong *oskar* mRNA mislocalisation phenotypes of *poulpe* mutants resemble those seen when the microtubule network is not correctly polarised, we examined the organisation of the microtubules by expressing kinesin-bgal, a constitutively active form of kinesin fused to b-galactosidase [27]. In wild-type ovaries, Kinesin-bgal localises to the posterior of the oocyte at stage 9 by moving along the weakly polarised microtubule network to the region with more plus ends than minus ends, just like *oskar* mRNA (Figure 1I) [23]. By contrast, Kinesin-bgal forms a cloud in the middle of most *poulpe* mutant oocytes, indicating that microtubule plus ends are concentrated in the centre (Figure 1M). Wild-type oocytes show an anterior-posterior gradient of microtubule density, with the strongest staining near the anterior/ lateral cortex, where the more stable, minus ends reside (Figure 1J). *poulpe* mutant oocytes, on the other hand, show high microtubule staining all around the oocyte cortex, with little signal in the centre, suggesting that microtubules are nucleated from the entire cortex (Figure 1N). These microtubule and *oskar* mRNA phenotypes of *poulpe* are very similar to those of strong *par-1* alleles, where ncMTOCs form all around the oocyte cortex and nucleate microtubules that extend towards the centre of the oocyte [18,24,28]. We therefore used a GFP protein trap insertion in Par-1 to examine whether it is recruited normally to the posterior cortex [20]. There is no obvious Par-1 enrichment at the posterior of *poulpe* mutant oocytes, however, indicating that polarity establishment is disrupted upstream of Par-1 localisation (Figure 1K and O). Consistent with this, the oocyte nucleus, which always moves to the anterior of wild-type and *par-1* mutant oocytes, is found in the middle of 10% of *poulpe* mutant stage 9 oocytes (n=20) (Figure 1L and P).

Recombination and deficiency mapping placed *poulpe* in the 400kb region between 84D14 and 84E8-9 and all three alleles failed to complement P{PZ}*unc-4503692*, a lethal P element insertion in the *unc-45* locus [29]. Sequencing revealed that *poulpe^6C3-11^* and *poulpe^4F2-4^* both have premature stop codons (Q250 -> Stop; Q 573 -> Stop) in the *unc-45* coding region and are therefore presumably null alleles, whereas the third allele *poulpe^4B4-10^* is likely to be a rearrangement. Thus, Poulpe corresponds to Unc-45, which is a TPR (tetratricopeptide repeat) and UCS (UNC-45, CRO1, She4p) domain containing protein that acts as a chaperone for folding and stabilising myosins [30, 31].

The weak phenotype of some *unc-45* germline clones resembles that of mutants in the single *Drosophila* myosin V, *didum*, in which Staufen and *oskar* mRNA are not anchored at the posterior cortex [32, 33]. *didum* mutants do not affect the localisation of Par-1, however, indicating that Unc-45 must be required for the function of another myosin that plays a role in polarity establishment. Although it is unclear how many of the 14 *Drosophila* myosins require Unc-45, many can be ruled out because their mutants are homozygous viable and fertile or because they are not expressed in the ovary [34]. We also excluded the Myosin VI, *jaguar*, as homozygous mutants have no effect on the posterior localisation of Staufen (Figure S2).

The most obvious candidate for a myosin involved in polarity establishment is non-muscle myosin II (MyoII), given its key role in the polarisation of the *C. elegans* zygote [35, 36]. MyoII is a hexamer formed by two molecules of the myosin heavy chain, Zipper (Zip), that dimerise through their long coiled coil tail domains and two copies of the essential light chain and the myosin regulatory light chain (MRLC), Spaghetti squash, which both bind to the neck region of each heavy chain [37] . To test whether Unc-45 is required for the folding and assembly of MyoII, we examined the distribution of endogenous Zipper, using a GFP protein trap insertion [38]. In wild-type egg chambers, Zipper is strongly enriched at the apical cortex of the follicle cells and localises at lower levels all around the oocyte cortex. Zippper GFP signal can be resolved from the apical follicle cell signal at high magnification at stage 10A and is most obvious at the nurse cell/oocyte boundary where there are no follicle cells (Figure 2A, A’). This cortical signal is lost in *unc-45* mutant germline clones and Zip-GFP is instead found in aggregates throught the nurse cell and oocyte cytoplasm, indicating that the formation of functional MyoII is impaired (Figure 2B, B’). MyoII performs many essential functions in the cell, including driving cytokinesis, and loss of function germline clones in the MRLC, *sqh*, therefore produce a range of defects, such as binucleate nurse cells and germline cysts with the wrong number of cells [39]. However, 64% of the mutant germline cysts that develop normally until stage 9 fail to localise Staufen to the posterior pole of the oocyte (n=47) (Figure 2C and D). This phenotype does not result from a defect in Staufen/*oskar* mRNA transport, which depends on microtubules rather than actin and is instead caused by a failure to establish anterior-posterior polarity, as shown by the loss of the posterior crescent of Par-1 (Figure 2E and F).

**Figure 2.**
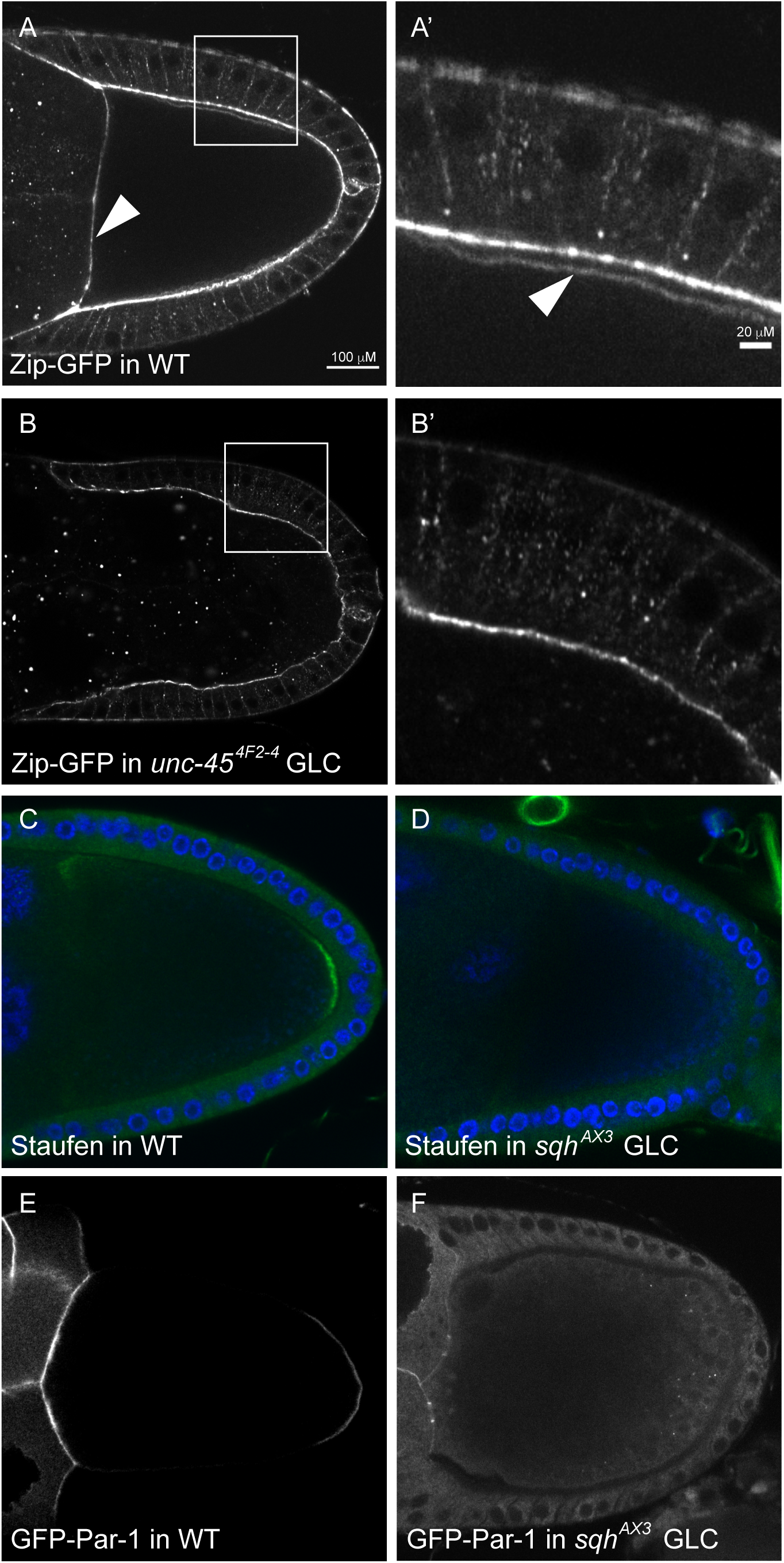
MyoII is chaperoned by Unc-45 and is required for the oocyte polarisation. (A and A’) A confocal image of a wild-type stage 9 egg chamber expressing Zipper-GFP, a protein trap line in the Myosin heavy chain. MyoII localises strongly to the apical side of the follicle cells and more weakly around the cortex of the oocyte beneath (arrowhead in A’). The arrowhead in A shows the anterior cortex of the oocyte where it abuts the nurse cells. (B and B’) A confocal image of an *unc-45*^4F2-4^ germline clone expressing Zipper-GFP. MyoII is lost from the oocyte cortex and accumulates in cytoplasmic aggregates, indicating that Unc45 is a MyoII chaperone. (C) Antibody staining of Staufen protein (green) in a wild-type oocyte, showing its localisation at the posterior cortex; DAPI (blue). (D) Antibody staining of Staufen protein (green) in a *sqh ^AX3^* mutant oocyte. Staufen is not localised posteriorly. (E) GFP-Par-1 expressed from a protein trap insertion localises at the posterior of a wild type oocyte. (F) GFP-Par-1 is not enriched at the posterior in *sqh^AX3^* mutant oocytes.

The requirement for MyoII in oocyte polarisation raises the question of whether MyoII itself is polarised. Live imaging of the Zipper GFP protein trap line reveals that MyoII is concentrated in a line at the posterior cortex of the oocyte, whereas the MyoII signal is more diffuse and weaker around the lateral cortex, suggesting that MyoII is asymmetrically activated at the posterior of the oocyte (Figure 3A, A’). An identical posterior enrichment is also observed for a Sqh-GFP transgene (Figure 3B, B’).

**Figure 3.**
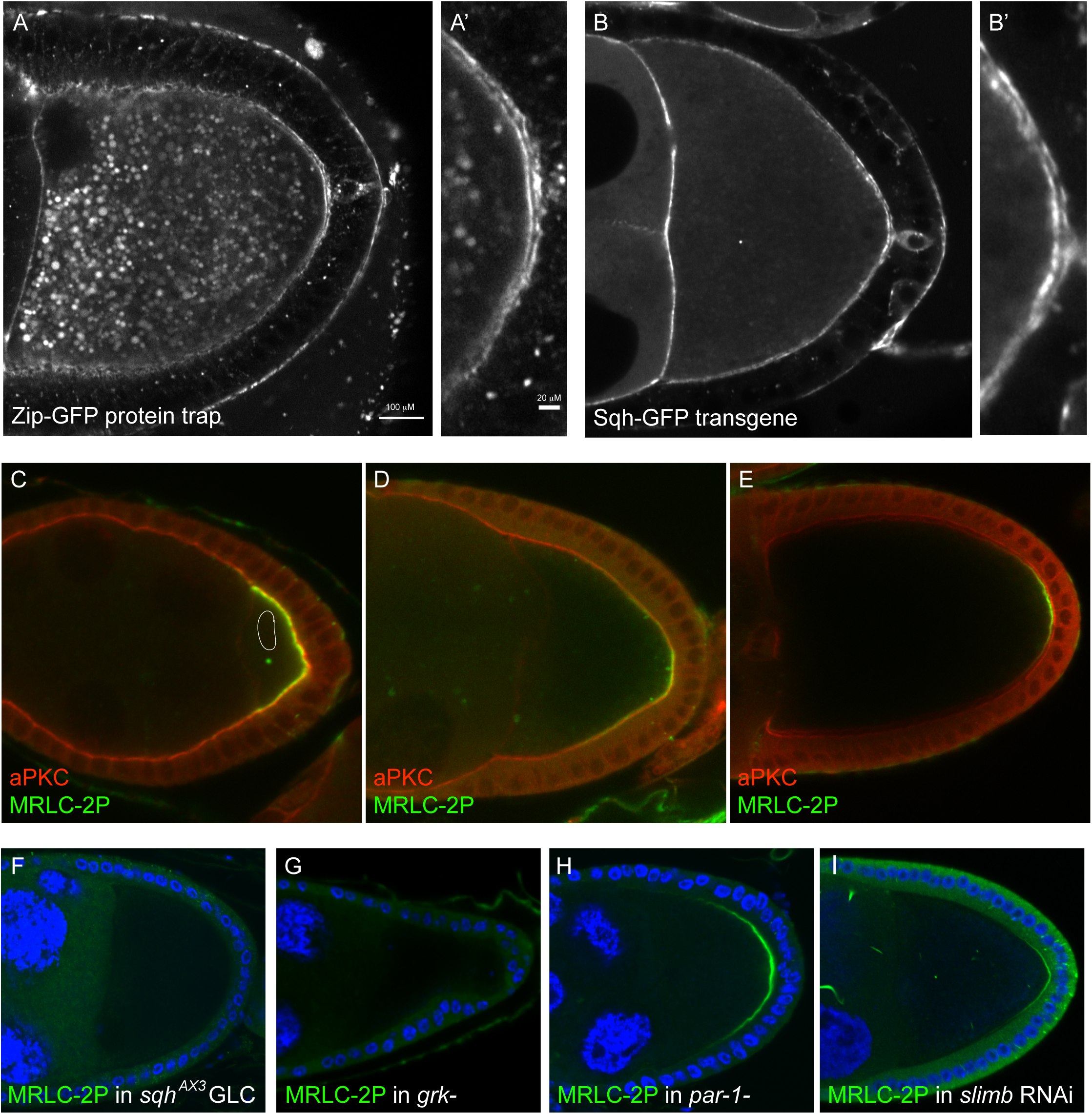
The myosin regulatory light chain is di-phosphorylated at the posterior cortex of the oocyte. (A and A’) A confocal image of a living egg chamber expressing Zipper-GFP (MyoII). MyoII is enriched at the posterior cortex of the oocyte. A’ shows a close-up of the posterior cortex. (B and B’) A confocal image of a living egg chamber expressing Sqh-GFP (MRLC). MRLC is enriched at the posterior cortex of the oocyte (close-up in B’). (C-E) Antibody staining for MRLC-2P (green) in stage 6 (C), stage 9 (D) and stage 10b (E) wild-type egg chambers. MRLC-2P localises to the regions of the oocyte cortex that contact the follicle cells at stage 6, becomes restricted to the posterior as the main body follicle cells migrate to cover the oocyte at stage 9 (D) and persists there until stage 10b (E). (F) MRLC-2P staining in a *sqh*^AX3^ germline clone. No MRLC-2P signal is detected, confirming the specificity of the antibody. (G) MRLC-2P staining in *grk^2B6/2E12^* mutant egg chamber. The posterior crescent of MRLC- 2P is lost, indicating that its formation depends on signalling from the posterior follicle cells. (H) MRLC-2P staining in a *par-1^6323/W3^* mutant egg chamber. The posterior crescent of MRLC-2P forms normally and maybe slightly expanded. (I) MRLC-2P staining in an egg chamber expressing in *slimb* RNAi in the germ line under the control of nos-Gal4. The posterior crescent of MRLC-2P forms normally in the absence of SCF/Slimb function.

MyoII activity is regulated by the phosphorylation of the conserved Threonine 20 and Serine 21 of MRLC, which activate its ATPase and motor activities [40, 41]. We therefore took advantage of specific antibodies that recognise *Drosophila* MRLC (Sqh) that is monophosphorylated on just Serine 21, the main activating site, or doubly phosphorylated on both Serine 21 and Threonine 20 (MRLC-2P) [42]. The monophosphorylated form of MRLC is enriched at the cortex but shows no obvious asymmetry along the anterior-posterior axis of the oocyte (Figure S3). By contrast, MRLC-2P is strongly enriched at the posterior cortex of the oocyte from stage 7 onwards (Figure 3C). This signal initially encompasses the entire region of the oocyte cortex that contacts the follicle cells, but as the main-body follicle cells surrounding the nurse cells migrate posteriorly to cover the oocyte during stage 9, MRLC-2P becomes restricted to a posterior crescent, where the signal persists until late stage 10B (Figure 3D, E). The localisation pattern of MRLC-2P therefore corresponds to the regions of the oocyte cortex that underlie the posterior terminal population of follicle cells, while the timing of the appearance of MRLC-2P coincides with the signal that polarises the oocyte.

To test whether MRLC di-phosphorylation depends on the polarising signal from the follicle cells, we examined MRLC-2P in various polarity mutants. As expected, no MRLC-2P is detected in *sqh^AX3^* null mutant germline clones, confirming that the signal is specific for phosphorylated MRLC (Figure 3F). More importantly, MRLC-2P is also completely lost from the posterior cortex of the oocyte in *gurken* mutants, which do not specify the posterior follicle cells and lack the polarising signal (Figure 3G). The posterior MRLC-2P crescent forms normally in *par-1* mutants, however, and may even expand, indicating that MRLC phosphorylation is upstream of Par-1 recruitment in the polarity signalling pathway (Figure 3H). The Slimb Ubiquitin ligase is the only known factor that acts upstream of Par-1 localisation in the oocyte except for the actin cytoskeleton [25]. Oocytes expressing Slimb RNAi still form the MRLC-2P posterior crescent (Figure 3I). Thus, MRLC is phosphorylated in response to the polarising signal and lies upstream of Slimb and Par-1 in the signal transduction pathway.

The discovery that MRLC is specifically di-phosphorylated at the oocyte posterior raises the question of whether this modification is required for oocyte polarity or is just a marker for this process. To test this, we took advantage of Sqh-GFP transgenes that cannot be di- phosphorylated because the minor phosphorylation site, Threonine 20, is mutated to Alanine (*sqh*AS) [39, 43]. Although the *sqh*-AS transgenes failed to rescue the polarity phenotype of *sqh^AX3^* null mutant germline clones, none of the available wild-type *sqh-*TS-GFP transgenes could rescue either, presumably because they are either not expressed at high enough levels or because the GFP tag impairs their function. We therefore generated new *sqh*TS and *sqh*AS transgenes without the GFP tag and under the endogenous *sqh* promoter and tested whether they had a dominant negative effect when present in two copies in a heterozygous *sqh*^AX3^/+ background. The wild-type *sqh*TS transgene had no effect on oocyte polarity as assayed by the posterior localisation of Staufen (Figure 4A, A’). By contrast, Staufen was not localised in half of the egg chambers over-expressing *sqh*AS, suggesting that the fully phosphorylated form of the MRLC plays an essential role in defining the posterior (Figure 4B,B’).

**Figure 4.**
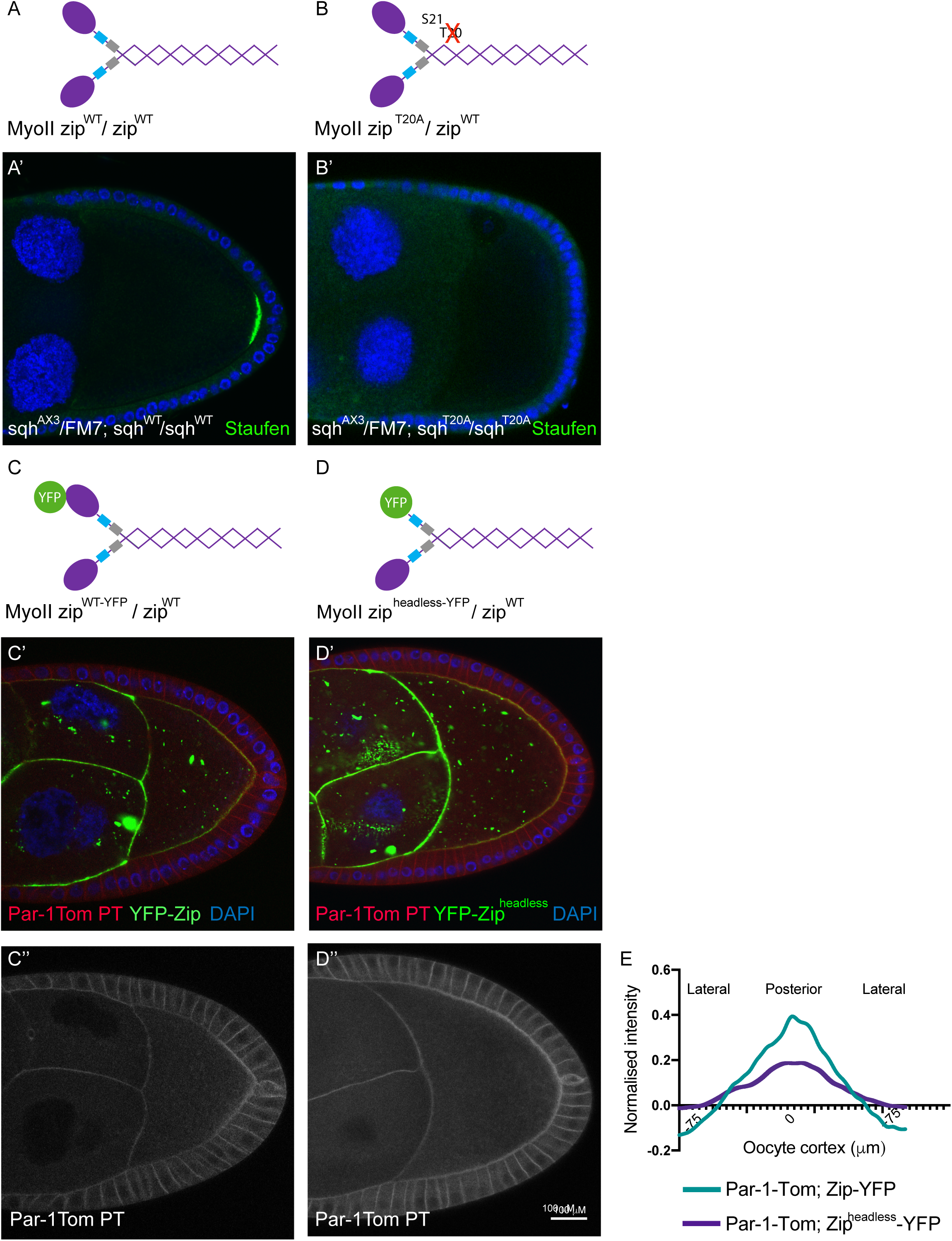
MRLC di-phosphorylation is required for anterior-posterior axis formation. (A) A diagram of the structure of wild type MyoII, with the Myosin Heavy Chain, Zipper, shown in purple, the Essential Light Chain in light blue and the Myosin Regulatory Light Chain (MRLC) in grey. (A’) Staufen staining (green) of a *sqh*^AX3^/FM7; *sqh*^WT^/*sqh*^WT^ egg chamber, expressing one endogenous copy and two transgenic copies of MRLC; DAPI (blue). Staufen localises normally to the posterior cortex of the oocyte in all cases. (B) A diagram of MyoII containing one copy of wild type MRLC and one copy of MRLC in which Threonine 20 is mutated to Alanine (T20A). (B’) Staufen staining of a *sqh*^AX3^/FM7; *sqh*^T20A^/*sqh*^T20A^ egg chamber, expressing one endogenous copy of wild-type MRLC and two transgenic copies of MRLC that cannot be phosphorylated on Threonine 20. Staufen fails to localise to the posterior in 48% of stage 9-10 oocytes of this genotype (n=152). (C) A diagram of wild type MyoII containing one copy of the Myosin heavy chain tagged by YFP. (C’ and C”) A YFP—Zipper/+ egg chamber showing the localisation of MyoII (green), Par- 1-Tomato (red) and DAPI (blue). (C”) shows Par-1-Tomato (white), which forms a crescent at the posterior of the oocyte and localises to the lateral cortex of the follicle cells. (D) A diagram of MyoII containing one copy of the Zipper in which the Myosin head has been deleted and replaced by YFP (YFP-Zipper^headless^). (D’ and D”) A YFP-Zipper^headless^/+ egg chamber showing the localisation of MyoII (green), Par-1-Tomato (red) and DAPI (blue). (D”) shows Par-1-Tomato (white), which forms a broad and weak crescent at the posterior of the oocyte. (E) Quantification of Par-1 Tomato localisation along the lateral and posterior oocyte cortex in YFP—Zipper/+ and YFP-Zipper^headless^/+ oocytes. The posterior pole, marked by the position of the polar follicle cells, lies at 0μm. The intensity was measured as the ratio of the PAR-1-Tomato signal at the oocyte cortex to the lateral signal in the follicle cells to normalise between different egg chambers.

The second phosphorylation of the MRLC on Threonine 20 has a little effect on the ATPase activity of MyoII in vitro in the presence of high concentrations of actin, compared to the form in which just Serine 21 is phosphorylated, but does increase ATPase activity when actin is limiting and enhances the speed at which MyoII can translocate F-actin [41]. Thus, the doubly phosphorylated form of MRLC may generate higher forces and/or faster contractions. To investigate whether MyoII activity is important for oocyte polarity, we examined the effects of over-expressing a headless Myosin heavy chain (Zip-YFP^headless^) that can still bind both light chains and form dimers with endogenous MyoII, but cannot exert force on actin [44]. Over-expression of wild-type Zip-YFP has no effect on the posterior recruitment of Par-1, whereas Zip-YFP^headless^ over-expression strongly reduces and broadens the Par-1 crescent (Figure 4 C-E). This suggests that cortical tension plays a role in Par-1 recruitment, although we cannot rule out the possibility that the headless myosin also disrupts filament formation.

In *C. elegans*, polarity is established by the contraction of the actomyosin cortex towards the anterior that localises the anterior PAR proteins by advection [35,36,45]. To test whether a similar mechanism operates in *Drosophila*, we imaged endogenous MyoII foci in the oocyte cortex using a Zipper-GFP protein trap line. Kymographs tracking the signal along the lateral and posterior cortex over time show that the MyoII forms foci that appear and disappear in way that is reminiscent of the pulsatile contractions observed in various *Drosophila* epithelial cells during morphogenesis [46–49]. Unlike these morphogenetic processes, the myosin foci at the cortex of the oocyte do not undergo large lateral movements, as shown by the nearly horizontal lines produced by each focus in the kymograph (Figure 5B and Figure S4). This suggests that the cortex is constrained, perhaps by connections through microvilli to the overlying follicle cells. More importantly, the MyoII foci are brighter and last longer at the posterior cortex than at the lateral cortex. Quantifying these data reveals that the MyoII foci at the posterior cortex persist for an average of 277 seconds, which is more than twice as long as the duration of the foci at the lateral cortex (125 seconds) (Figure 5C). Thus, the di-phosphophorylation of Sqh increases the duration of actomyosin pulses, presumably leading to higher cortical tension.

**Figure 5.**
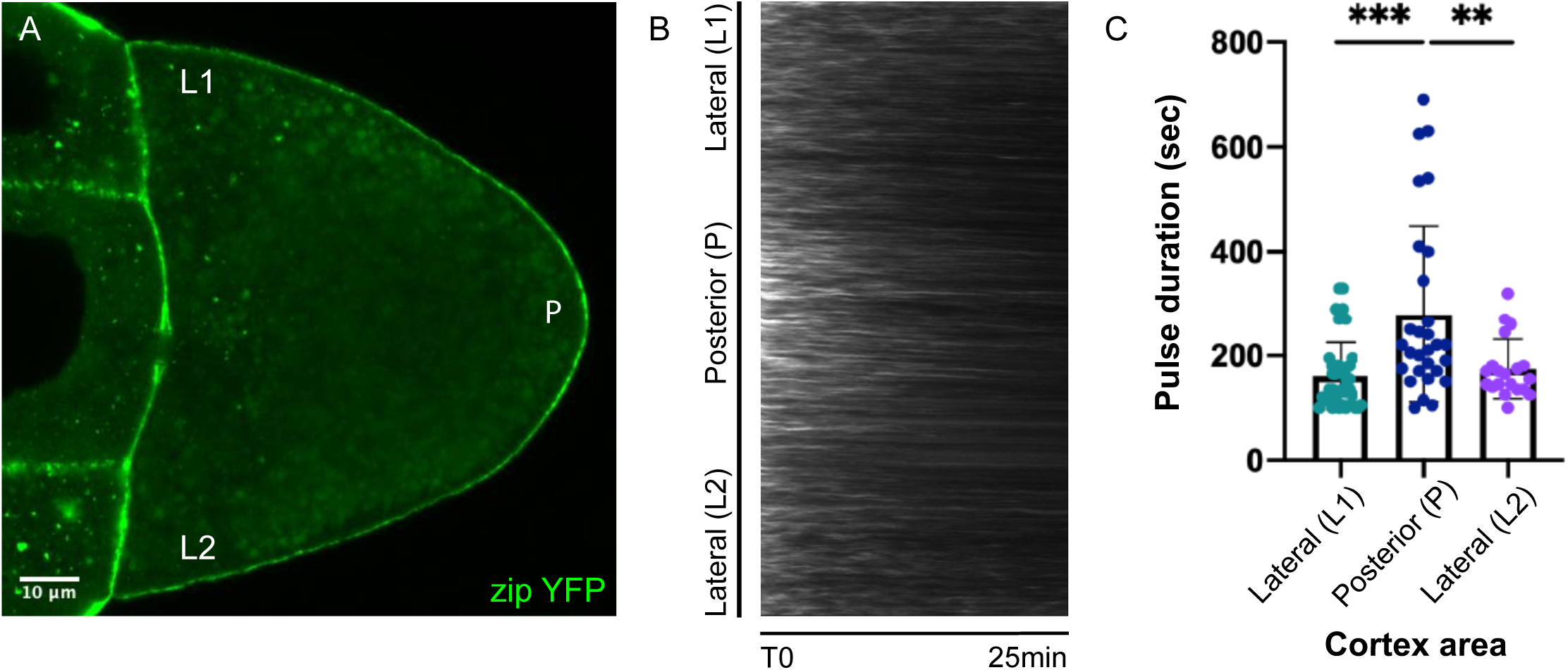
MyoII forms cortical foci that persist longer at the posterior. (A) A still image from a movie of wild-type egg chamber expressing Zipper-GFP in the germline. (B) A kymograph showing the changes in Zipper-GFP levels over time along the oocyte cortex. Zipper-GFP foci remain stationary, indicating that there is no cortical contraction, and oscillate in intensity over time. (C) A graph showing the durations of Zipper-GFP pulses at the lateral and posterior cortex (L1, P, L2). Pulse durations are measured using an automated detection and segmentation algorithm. The pulses at the posterior last twice as long as the lateral pulses.

The polarising cortical contraction in *C. elegans* is a single, transient event that occurs in response to sperm entry early in the first cell cycle. There is no clear morphological sign that indicates when the signal to polarise the *Drosophila* oocyte is produced, however, and we therefore cannot exclude the possibility that there is a cortical contraction that we have not succeeded in visualising sometime during the 12 or more hours between stages 6-9. If this is the case, MyoII activation should be transiently required to establish polarity but would not be needed to maintain it once the Par proteins are asymmetrically localised. To test this, we examined the effects of acutely inhibiting MRLC kinases after the posterior Par-1 crescent has formed. In many contexts, MyoII is activated by the Rho-dependent kinase, Rok, which is inhibited by Y-27632 [50, 51]. However, treating egg chambers with Y-27632 has no effect on posterior Par-1 recruitment or myosin phosphorylation (Figure 6A, B and G). Consistent with this, Rho activity, as measured by the AniRBD-GFP reporter, is lower at the posterior cortex of the oocyte than elsewhere (Figure 6E) [52]. Furthermore, treatment with higher concentrations of Y-27632 causes an expansion of the posterior Par-1 crescent rather than a loss, presumably because these concentrations also inhibit aPKC, which phosphorylates Par-1 to exclude it from the lateral cortex {Atwood:2009cm, Doerflinger:2006bi) (Figure 6C). This confirms that Y-27632 enters the oocyte efficiently and is active, ruling out Rok as the kinase that phosphorylates Sqh at the posterior. By contrast, exposing egg chambers to ML-7, an inhibitor of myosin light chain kinase, leads to a complete loss of posterior Par-1 and Sqh-2P in 15 minutes (Figure 6D, F and H). This confirms that MyoII phosphorylation is required to localise Par-1 at the posterior and indicates that this is continuous requirement, ruling out a contraction-based mechanism for polarity establishment.

**Figure 6.**
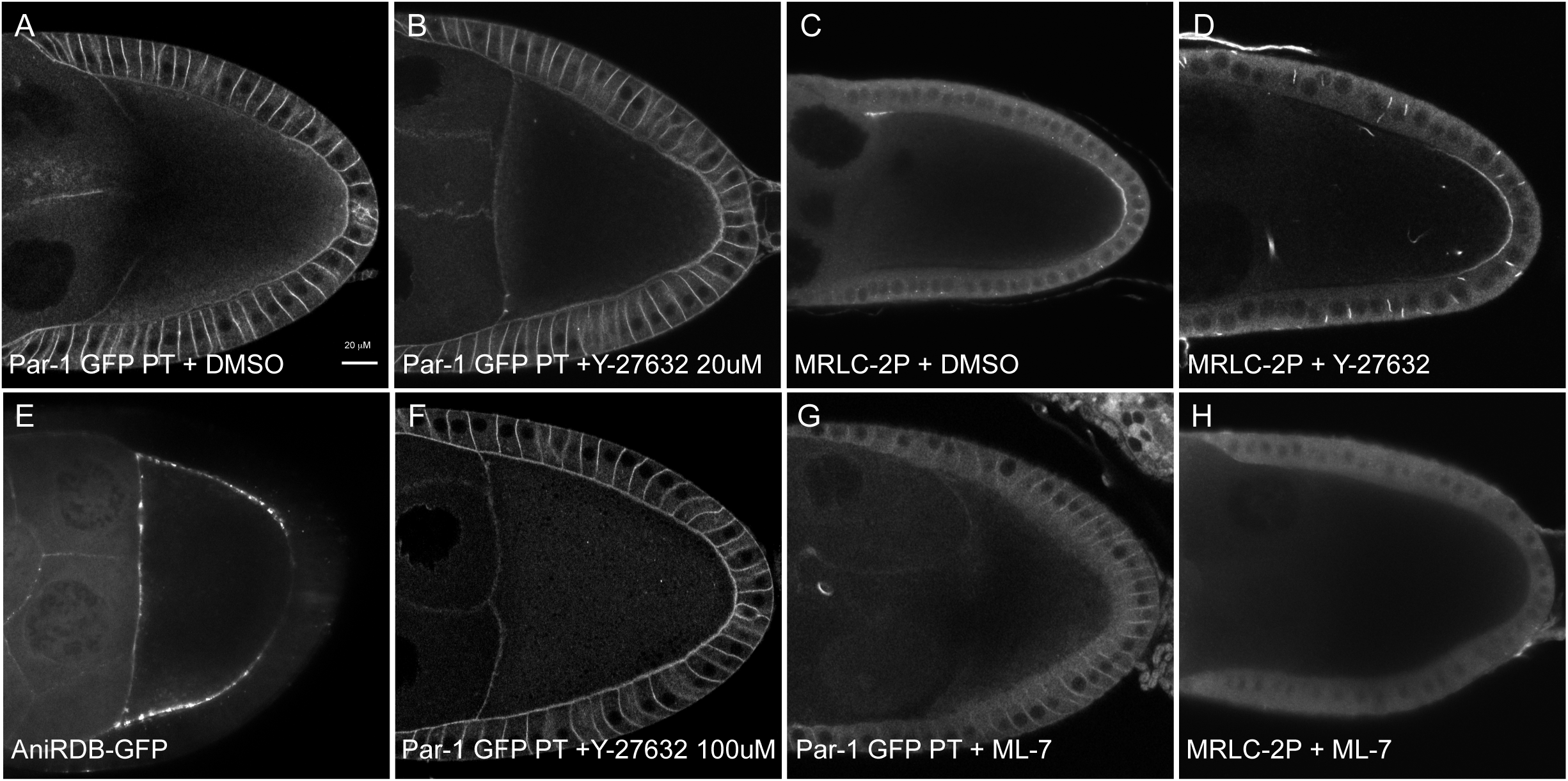
MyoII activation is inhibited by the MLCK inhibitor ML-7. (A) A confocal image of an egg chamber expressing Par-1 GFP from a protein trap insertion after incubation in DMSO. Par-1 GFP localises in a crescent at the posterior cortex of the oocyte. (B) A confocal image of egg chambers expressing Par-1 GFP after incubation in the Rok kinase inhibitor Y-27632 20μM. Y-27632 has no effect on Par-1 localisation. (C) A confocal image of MRLC-2P immunostaining in wild type egg chambers after incubation in DMSO. MRLC-2P signal forms a crescent at the posterior cortex of the oocyte. (D) A confocal image of MRLC-2P immunostaining in wild type egg chambers after incubation in the Rok kinase inhibitor Y-27632 20μM. Y-27632 has no effect on MRLC-2P localisation. (E) A confocal image of an egg chamber expressing the AniRBD-GFP reporter for active Rho-GTP. The AniRBD-GFP signal is lower at the posterior cortex of the oocyte than elsewhere. (F) A confocal image of egg chambers expressing Par-1 GFP after incubation in the Rok kinase inhibitor Y-27632 100μM. At this high concentration, Par-1 GFP expands around the lateral cortex of the oocyte, presumably because this concentration inhibits aPKC. (G) A confocal image of an egg chamber expressing Par-1 GFP oocyte after incubation in the myosin light chain kinase inhibitor ML-7. Par-1-GFP no longer localises at the posterior cortex of the oocyte. (H) A confocal image of MRLC-2P immunostaining in wild type egg chambers after incubation in the myosin light chain kinase inhibitor ML-7. MRLC-2P signal no longer localises at the posterior cortex of the oocyte.

**Figure 7.**
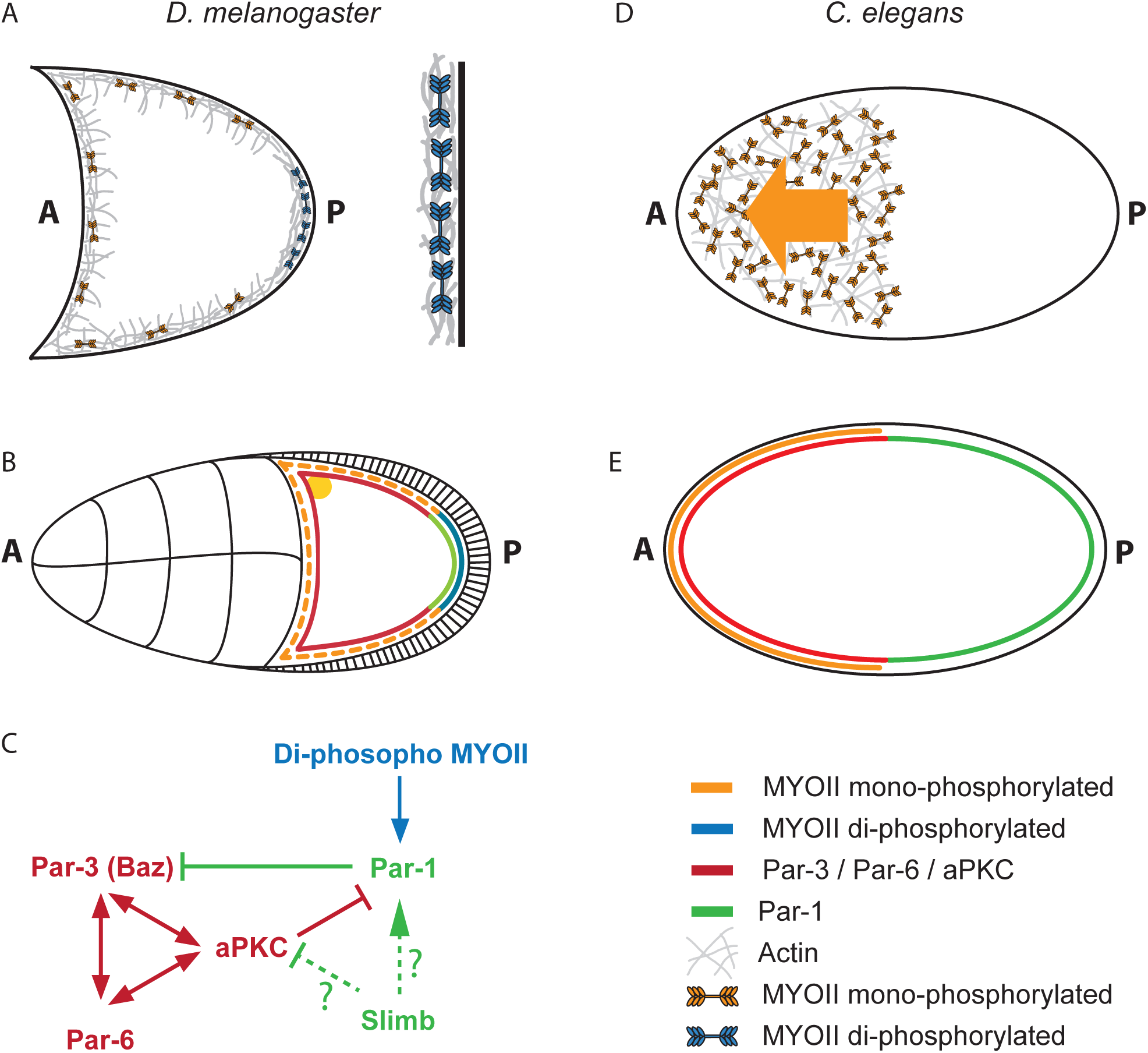
Model for the role of MyoII in the *Drosophila* oocyte polarisation. (A) Mono-phosphorylated MyoII is distributed all along the cortex of the *Drosophila* oocyte whereas di-phosphorylated MyoII concentrates at the posterior. Phosphorylated MyoII hexamers form bipolar filaments and generate force between two anti-parallel actin filaments, increasing cortical tension. (Mono-phosphorylated MyoII: orange, Di-phosphorylated MyoII: blue, Actin (grey). (B) MyoII localises all along the oocyte cortex, but is specifically di-phosphorylated along the posterior of the oocyte from stage 6 (blue), leading to the posterior recruitment of Par-1 (green) and the exclusion of aPKC/Par-3/Par-6(red). (Mono-phosphorylated MyoII: orange, Di-phosphorylated MyoII: blue,, Actin: grey). (C) In the *C.elegans* zygote, MyoII drives a contraction of the actomyosin cortex towards the anterior, which moves the anterior PAR proteins with it by advection, leading to the asymmetric distribution of PAR proteins and the establishment of AP polarity. (MyoD: orange, Actin: grey) (D) The anterior contraction of MyoII in the *C. elegans* zygote results in the asymmetric distribution of aPKC, Par-3 and Par-6 along the anterior cortex and Par-2 and Par-1 along the posterior cortex (MyoII: orange, Par-1: green, aPKC/Par-3/Par-6: red). (E) Proposed signalling pathway for the establishment of oocyte AP polarity. Slimb may act to promote the posterior recruitment of Par-1 in response to the diphosphorylation of MyoII or could act to exclude the anterior Par proteins.

## Discussion

Although it was discovered more than twenty years ago that the posterior follicle cells signal to polarise the anterior-posterior axis of the oocyte, almost nothing is known about the nature of this signal or how it is transduced to the oocyte. Here we show that a key response to the signal is the di-phosphorylation of MRLC at the posterior of the oocyte, as the appearance of MRLC-2P coincides with where and when the polarising signal is produced and depends on the specification of the posterior follicle cells by Gurken. More importantly, MRLC di-phosphorylation is required for all subsequent steps in oocyte polarisation, since a form of MRLC that can only be mono-phosphorylated on Serine 21 acts as a dominant negative to disrupt axis formation and inhibiting the phosphorylation prevents the recruitment of Par-1 to the posterior cortex of the oocyte.

MRLC-2P shows a very different distribution from MRLC-1P in both *Drosophila* and Ascidian embryo morphogenesis, but its function in vivo has remained unclear (Sherrard et al, 2010; Zhang and Ward, 2011). Our results therefore provide the first example where the di-phosphorylation of MRLC has been demonstrated to play an essential role in development. Vasquez et al. found that the second phosphorylation of MRLC on Threonine 20 has a negligible effect on MyoII’s ATPase activity in vitro, but causes a decrease in the rate of actin translocation and in the rate of apical constriction in the mesoderm of the gastrulating embryo, suggesting that this modification increases the force generated by MyoII [41]. Because of the clear spatial distribution of MRLC-2P in the oocyte cortex, our analysis reveals a second effect of the phosphorylation of the Threonine 20, which is that it more than doubles the duration of MyoII pulses. This may simply reflect an increase in the time that it takes Myosin phosphatase to remove two phosphates, instead of one, or be due to a more complicated effect on the structure of the myosin hexamer. Nevertheless, this second phosphate allows MyoII to generate more force for longer than the mono-phosphorylated form.

The critical function of MRLC-2P in the oocyte is to trigger the recruitment of Par-1 to the posterior cortex, raising the question of how this occurs. The second phosphorylation of MRLC is likely to increase the force generated by MyoII and one would therefore expect to see higher contractility in the posterior oocyte cortex. However, in contrast to the mesoderm, where MRLC-2P increases contraction rates, this does not occur in the oocyte cortex, as there is no lateral movement of MyoII at the posterior or elsewhere. This may be because the actin cortex is different from the mesoderm and cannot contract, possibly because it is denser and rigidly anchored in place through the microvilli that connect to microvilli in the follicle cells. If this is the case, the extra force exerted by MyoII at the posterior should increase the stress on the cortex and on MyoII itself, and this may be the critical change that recruits Par-1 to the posterior. For example, MyoII or some other cortical component could act as a tension sensor that exposes a binding site for Par-1, similar to way in which Talin and a-catenin expose binding sites for Vinculin when stretched [53, 54]. This model can explain why the over-expression of headless Zipper disrupts Par-1 localisation, since this should result in mixed MyoII hexamers with fewer heads that therefore exert less force.

Any model for Par-1 recruitment must explain the role of Slimb in this process [25]. Par-1 and the anterior Par-3 (Baz)/Par-6/aPKC complex are mutually antagonistic, with Par-1 excluding Baz from the cortex by phosphorylation and aPKC excluding Par-1 by phosphorylation [21, 22]. The levels of the anterior polarity factors aPKC and Par-6 are increased in *slimb* mutants, leading to the suggestion that the Slimb/SCF Ubiquitin ligase normally targets them for degradation at the posterior, thereby allowing Par-1 to localise there. Thus MRLC-2P and the increased tension may promote the SCF-dependent removal of the anterior Par proteins from the posterior. However, it is also possible that Slimb/SCF plays an indirect role in polarity by reducing Par-6/aPKC levels everywhere and thereby setting a threshold for cortical exclusion of Par-1 by aPKC that is overcome specifically at the posterior by its MRLC-2P-dependent recruitment. Alternatively, Slimb may play a role that is independent of its regulation of aPKC and Par-6. It has recently been shown that the SCF Ubiquitin ligase complex regulates the ubiquitylation of Zipper in *Drosophila* auditory organs, while Par-1 contains a Ubiquitin Binding Associated (UBA) domain, raising the possibility that SCF ubiquitylates Zipper in response to tension to create a binding site for Par-1 [55]. In this context, it is worth noting that the *C elegans* myosin II heavy chain was first identified in an expression screen for proteins that bind to Par-1 and co-immuno- precipitates with Par-1 from embryos [56].

In both *Drosophila* and *C. elegans*, anterior-posterior axis is defined by the formation of complementary cortical domains of mutually antagonistic Par complexes. Our results reveal a further similarity in that Myosin activity is required to establish these PAR domains in each case. However, polarity in the worm is established by a myosin-driven contraction of the cortex towards the anterior that localises the anterior PAR proteins, whereas MyoII activation in the *Drosophila* oocyte localises Par-1 to the posterior. A second key difference between polarity establishment in worms and flies is that myosin activation is continuously required for Par-1 localisation in *Drosophila*, since this localisation is abolished by ML-7 treatment in oocytes that have already polarised. By contrast, the requirement for MyoII is transient in *C. elegans*, and MyoII contractility is not required to maintain PAR polarity once it is established [5]. This may reflect the different nature of the polarising cues in each system, since sperm entry in the worm is a one-off event, whereas the posterior follicle cells remain adjacent to the posterior of the *Drosophila* oocyte throughout oogenesis and are therefore in a position to provide the polarising signal continuously. These differences highlight the context-dependent relationship between the actomyosin cortex and polarity factors.

While the role of MyoII in polarity establishment in flies and worms is very different, there is a striking parallel between MyoII’s role in localising Par-1 posteriorly in the *Drosophila* oocyte and in localising the cell fate determinant Miranda basally during the asymmetric cell divisions of the neuroblasts. Like Par-1, Miranda is excluded from the cortex by aPKC phosphorylation and this was initially thought to be sufficient to explain its basal localisation in the neuroblast [57, 58]. It has recently emerged, however, that aPKC’s main function is to exclude Miranda from the apical and lateral cortex during interphase and that activated MyoII then recruits Miranda basally during mitosis in a process that is inhibited by ML-7 [59, 60]. Thus, Miranda and Par-1 appear to share a common localisation mechanism, which may provide a more general paradigm for the role of MyoII in generating cellular asymmetries.

## MATERIAL AND METHODS

### Key Resources Table

| REAGENT or RESOURCE | SOURCE | IDENTIFIER |
| --- | --- | --- |
| **Antibodies** |  |  |
| FITC-coupled anti-α-tubulin (1/200) | Sigma | F2168 |
| Staufen (1/200) | {Johnston:1991wj} | NA |
| SQH2P/MRLC-2P (1/100) | Zhang and Ward, 2011 | NA |
| SQH1P/MRLC-1P (1/100) | Zhang and Ward, 2011 | NA |
| SQH2P/MRLC-2P (1/100) | This study. Same peptides as Zhang and Ward, 2011 | Pocono Rabbit Farm & Laboratory |
| Alexa conjugated secondary antibodies (1/300) | Thermo Fisher Scientific | NA |
| **Bacterial Strains** |  |  |
| Bioline silver competent cells | BIOLINE UK LTD | BIO85026 |
| **Chemicals** |  |  |
| Wheat germ agglutinin, Texas Red®-X (1/300) | Invitrogen | W21405 |
| Texas Red™-X Phalloidin (1/300) | Thermo Fisher Scientific | T7471 |
| VectaShield Mounting Medium with DAPI | Vector Laboratories Ltd | H-1200-10 |
| Myosin light chain kinase inhibitor ML-7 | Merck Ltd | 475880 |
| ROCK inhibitor Y-27632 | Hello BIO | HB2297 |
| Schneider's Insect medium | Sigma Aldrich Company Ltd | S0146 |
| Insect Cell screened FBS | Thermo Fisher Scientific | SH30070.03I |
| BSA, fatty acid free | Sigma Aldrich Company Ltd | A4612-1G |
| Insulin Solution, Human | Sigma Aldrich Company Ltd | I9278-5ML |
| 16% formaldehyde methanol free | Fisher Scientific Worldwide | 10751395 |
| Tween 20 | Sigma | P1379 |
| **Critical Commercial Assays** |  |  |
| QuikChange II XL Site-Directed Mutagenesis kit | Agilent Technologies | 200521 |
| Gibson Assembly Master mix | NEW ENGLAND BIOLABS | E2611S |
| **Fly stocks** |  |  |
| See table 1 | NA | NA |
| **Oligonucleotides** |  |  |
| See table 2 | NA | NA |
| **Recombinant DNA** |  |  |
| pattB cloning vector | {Potter:2010eu} | NA |
| pattB -sqhWT | This study | NA |
| pattB -sqhT20A | This study | NA |
| **Software and Algorithms** |  |  |
| Olympus Fluoview Version 3.1. | NA | NA |
| Fiji Image Processing Software | {Schindelin:2012ir} | https://imagej.net/Fiji |
| Excel | Microsoft Office | NA |
| Prism 8 | Graphpad | NA |

### Stock maintenance and *Drosophila* genetics

Standard procedures were used for *Drosophila* husbandry and experiments. Flies were reared on standard fly food supplemented with dried yeast at 25 °C. Heat shocks were performed at 37 °C for 1 h (twice daily) for three days. Flies were kept at 25 °C for at least 3 to 5 days after the last heat shock before dissection. UAS-transgenes were expressed using Gal-4 drivers in flies raised at 25°C; adult females were dissected at least 3 to 5 days after they had eclosed.

### Drug treatment

Ovaries were incubated in a Schneider’s insect medium solution, 10% FBS and Insulin (1/2000) for 20 min with 20 μM ROCK inhibitor Y-27632 (HelloBio HB2297) or 100 uM myosin light chain kinase inhibitor ML-7 (Merck 475880) and fixed for 20 min in 4% paraformaldehyde and 0.2% Tween 20 in PBS.

The reversibility of ML-7 was tested by incubating the ovaries in a Schneider’s insect medium solution, 10% FBS and Insulin (1/2000) for 20 minutes with 100μM ML-7, followed by a 15 min wash in the Schneider’s insect medium and fixed for 20 min in 4% paraformaldehyde and 0.2% Tween 20 in PBS.

### Immunostaining

Ovaries were fixed for 15 min in 4% formaldehyde and 0.2% Tween 20 in PBS. For phospho-specific antibody immunostainings, a phosphatase inhibitor solution was added to the PBS 0,2% Tween 20 solution. 50X phosphatase inhibitor solution kept at -80°C: 0.105g NaF (Sigma S79209), 0.540g B glycerophosphate (Sigma G9422), 0.092g Na_3_VO_4_ (Sigma 450243), 5.579g Sodium pyrophosphate decahydrate (Sigma S6422), qsp 50 ml dH2O.

*α*-tubulin immunostainings: ovaries were fixed 10 min in 10% formaldehyde and 0.2% Tween 20 in PBS as described by Theurkauf et al. (1992).

Ovaries were then blocked in 10% bovine serum albumin (in PBS with 0.2% Tween 20) for at least 1 h at room temperature. Samples were incubated with primary antibodies for at least 3 h in PBS with 0.2% Tween 20 and 10% BSA and were then washed three times in PBS-0.2% Tween 20 for 30 min. They were then incubated in secondary antibodies for at least 3 h in in PBS-0.2% Tween 20 and washed again at least 3 times before mounting in Vectashield containing DAPI (Vector laboratories) The concentrations of primary antibodies used are indicated in the Key Resources Table. Secondary antibodies and Phalloidin were used at 1/300. Incubations with Wheat germ agglutinin (1/300) were performed in PBS with 0.2% Tween 20 for 30 min followed by a 30 min wash.

### In situ hybridisations

In situ hybridisations were carried out using RNA probes labelled with Digoxigenin-UTP [61].

### Imaging

Imaging was performed using an Olympus IX81 (40×/1.3 UPlan FLN Oil or 60×/1.35 UPlanSApo Oil). Images were collected with Olympus Fluoview Ver 3.1. Image processing was performed using Fiji (Schindelin et al., 2012).

### Molecular biology

To generate the pattB -sqhWT construct, 2.7 kb of sqh genomic DNA was amplified by PCR with the oligos H472 and H473 (see Table 2) and inserted in the PhiC31 integration pattB cloning vector (Potter et al., 2010) digested with XbaI-BamHI using the Gibson assembly method (Gibson Assembly Master mix NEB). To generate the pattB -sqhT20A construct, we used the Q5 Site Directed Mutagenesis kit (NEB) to generate the sqhT20A mutation in the pattB -sqhWT construct using oligos H334 and H335 (See Table 2). The pattB-sqhWT and pattB-sqhT20 constructs were injected into y v;;attP2 line *Drosophila* embryos [62] to generate transgenic lines.

**Table 1.** *Drosophila* mutant stocks and transgenic lines

| Genotype | Source | Identifier |
| --- | --- | --- |
| **Mutant alleles** |  |  |
| y<sup>1</sup> w<sup>1</sup> (used as wild type) | Bloomington Drosophila Stock Centre | 1495 |
| w ;; FRT82B unc-45<sup>4B4-10</sup>/TM3 | Martin et al. (2003). Called poulpe<sup>4B4-10</sup> | NA |
| w ;; FRT82B unc-45<sup>4F2-4</sup>/TM3 | Martin et al. (2003). Called poulpe<sup>4F2-4</sup> | NA |
| w ;; FRT82B unc-45<sup>6C3-11</sup>/TM3 | Martin et al. (2003). Called poulpe<sup>6C3-11</sup> | NA |
| gurken<sup>2B6</sup> b pr cn sca/CyO | Neuman-Silberberg et al, 1993 | NA |
| gurken<sup>2E12</sup> b / CyO | Neuman-Silberberg et al, 1993 | NA |
| w ; par-1<sup>W3</sup>/ CyO | Shulman et al., 2000 | NA |
| w ; par-1<sup>6323</sup>/CyO | Shulman et al., 2000 | NA |
| w FRT 19A sqh<sup>AX3</sup>/FM7 | Bloomington Drosophila Stock Centre | 25712 |
| Df(3R)jaguar<sup>322</sup>/TM3 | Bloomington Drosophila Stock Centre | 8776 |
| w ; FRTG13 didum 2R-234-17-3 / CyO | {Luschnig:2004wx}. Called shorty L744 | NA |
| **Tagged and/or mutant genes** |  |  |
| y<sup>1</sup> sc* v<sup>1</sup> sev<sup>21</sup>; slimb RNAi attP | Bloomington Drosophila Stock Centre | 33986 |
| w; UAS-zip-GFP | Bloomington Drosophila Stock Centre | 80156 |
| w; UAS-zip<sup>ΔN</sup> headless-YFP | Dawes-Hoang et al., 2005 | NA |
| y w; Pin / CyO; Kinesin:βgal | Clark et al., 1994 | KZ503 |
| w ; mat-tub-GFP-par-1/ CyO | Huynh et al., 2001 | NA |
| w ; UASp-GFP:par-1(N1S)-GFP/CyO | Huynh et al., 2001 | NA |
| par-1-GFP protein trap / CyO | {Lighthouse:2008cd} | NA |
| par-1-Tomato protein trap / CyO | This study | NA |
| w ; zip-GFP protein trap insertion | Kyoto stock centre | 115082 |
| y<sup>1</sup> w* cv<sup>1</sup> sqhAX3; sqh-GFP | Bloomington Drosophila Stock Centre | 57144 |
| w ; sqh<sup>WT</sup> attP2 | This study | NA |
| w ; sqh<sup>T20A</sup> attP2 | This study | NA |
| w ; P{tdTomato-0}N1001.5a | St Johnston lab | NA |
| **Gal4 drivers** |  |  |
| w ; mat-tub:Gal4 | St Johnston lab and Jean-Paul Vincent | NA |
| w ; mat-tub:Gal4-VP16 | St Johnston lab and Jean-Paul Vincent | 7063 |
| w ; nanos:Gal4 VP16 | {VanDoren:1998in} | NA |
| **FRT lines** |  |  |
| w ;; FRT82B ubi-GFP | Bloomington Drosophila Stock Centre | 5188 |
| w ;; FRT 82B ovoD / TM3 | Bloomington Drosophila Stock Centre | 2149 |
| y w hs:flp FRT 19A ovoD / <u>C(1)DX</u> | Bloomington Drosophila Stock Centre | 23880 |
| FRT 19A GFP ; hs:flp | St Johnston lab | NA |
| **Landing site line** |  |  |
| y v ;; attP2 | Bloomington Drosophila Stock Centre | 31207 |
| **Transposase line** |  |  |
| w* ; Sc/CyO [Hop] | St Johnston lab | NA |

**Table 2.**
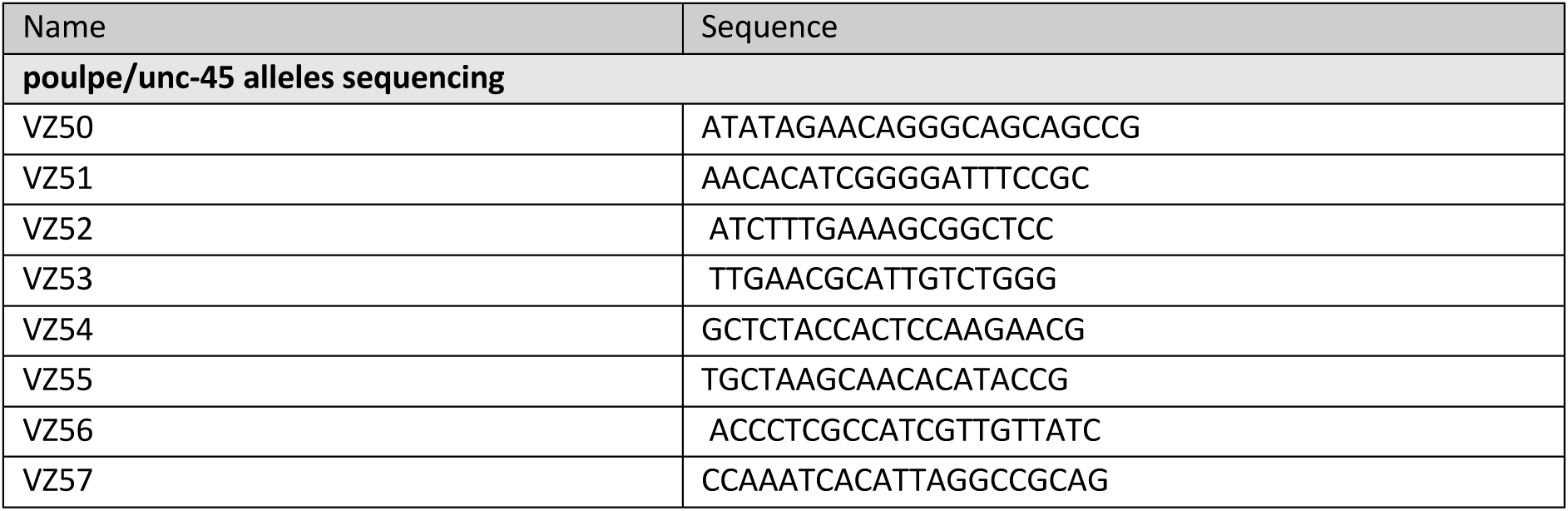
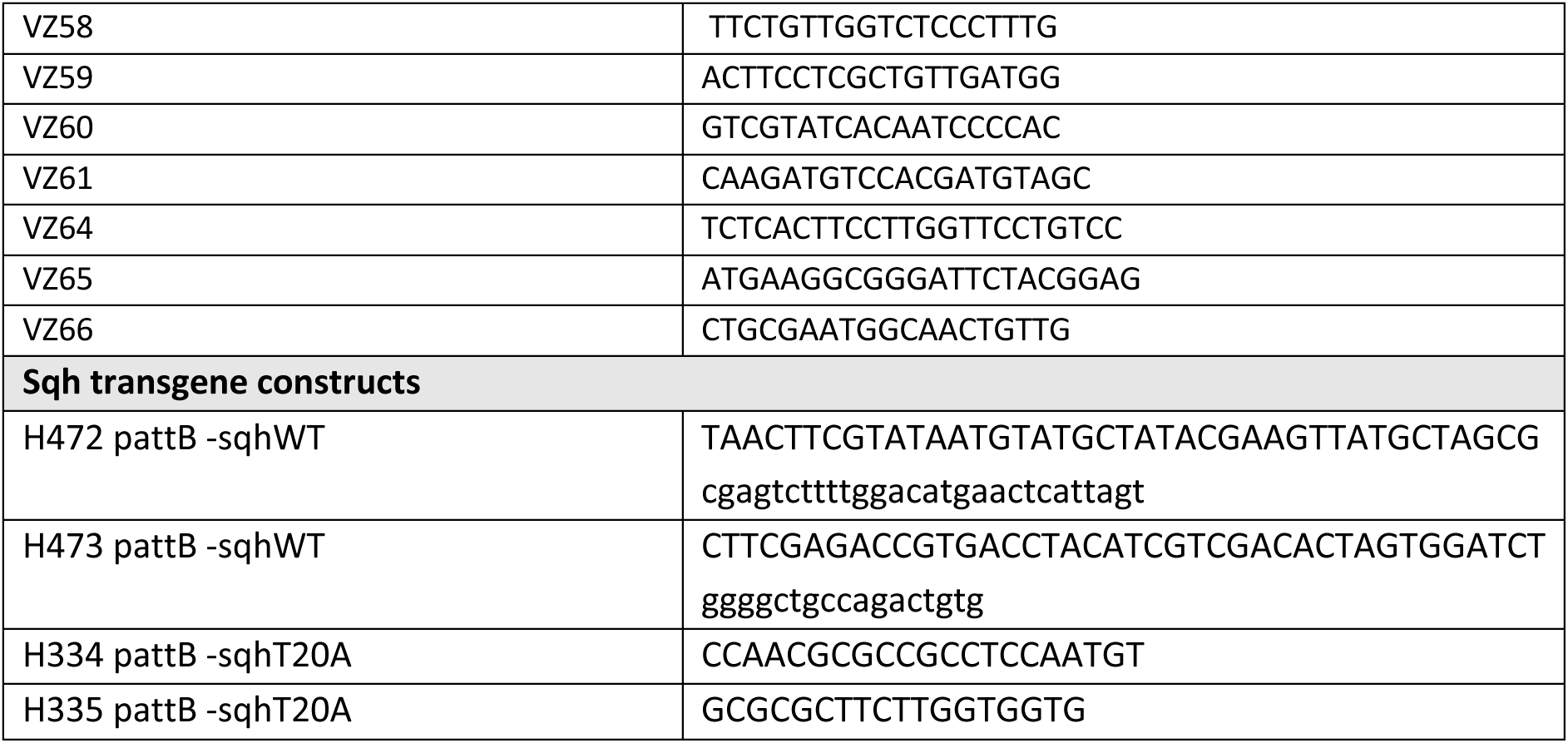
Oligonucleotides

The par-1-Tomato protein trap was generated by replacing the GFP tag of the par-1-GFP protein trap by the Tomato tag using the P swap technique [63]. A Tomato transposon donor line in the appropriate reading frame (PIGP3{tdTomato-1}) was crossed with the Par- 1-GFP protein line together with the Hop transposase. Larvae from this cross were screened with a Leica MZ16 fluorescent microscope and individual red fluorescent larvae were picked into a fresh vial. Adults were crossed to CyO for balancing.

### Analysis of MyoII pulses

The distribution of MyoII along the oocyte cortex was imaged by recording the fluorescence from a UAS-Zipper-GFP line expressed in the germline. The flies were dissected under Voltalef 10S oil and imaged at 40x magnification on a confocal microscope. The Zipper-GFP signal was imaged with a 40x 1.3 NA objective once every 15 sec for 25 min with a pixel size of 0.198 µm. The kymograph was generated using Fiji (Schindelin et al., 2012), the quantification of NMYII pulse times was perf with Fiji. The durations of MyoII pulses signal were automatically measured by tracking adjacent strong intensity pixels in the kymograph. 25 consecutive measurements were pooled to determine the average time of NMYII expression at both lateral sides and at the posterior of the oocyte cortex.

## Acknowledgements

We would like to thank Richard Ward for sharing the SQH1P and SQH2P antibodies, Andrea Brand for the zip^ΔN headless^-YFP and Richard Butler at the Gurdon Institute Imaging Facility for help with image analysis and quantification. This work was supported by a Wellcome Trust Principal Fellowship to D St J (080007, 207496) and by centre grant support from the Wellcome Trust (092096, 203144) and Cancer Research UK (A14492, A24823).

## Author Contributions

The project was conceived and designed by all authors. H.D. and V. Z. performed the experiments and the data analysis. D. St J. wrote the manuscript, which was edited by H. D. and V. Z.

---

**Supplementary figure 1.**
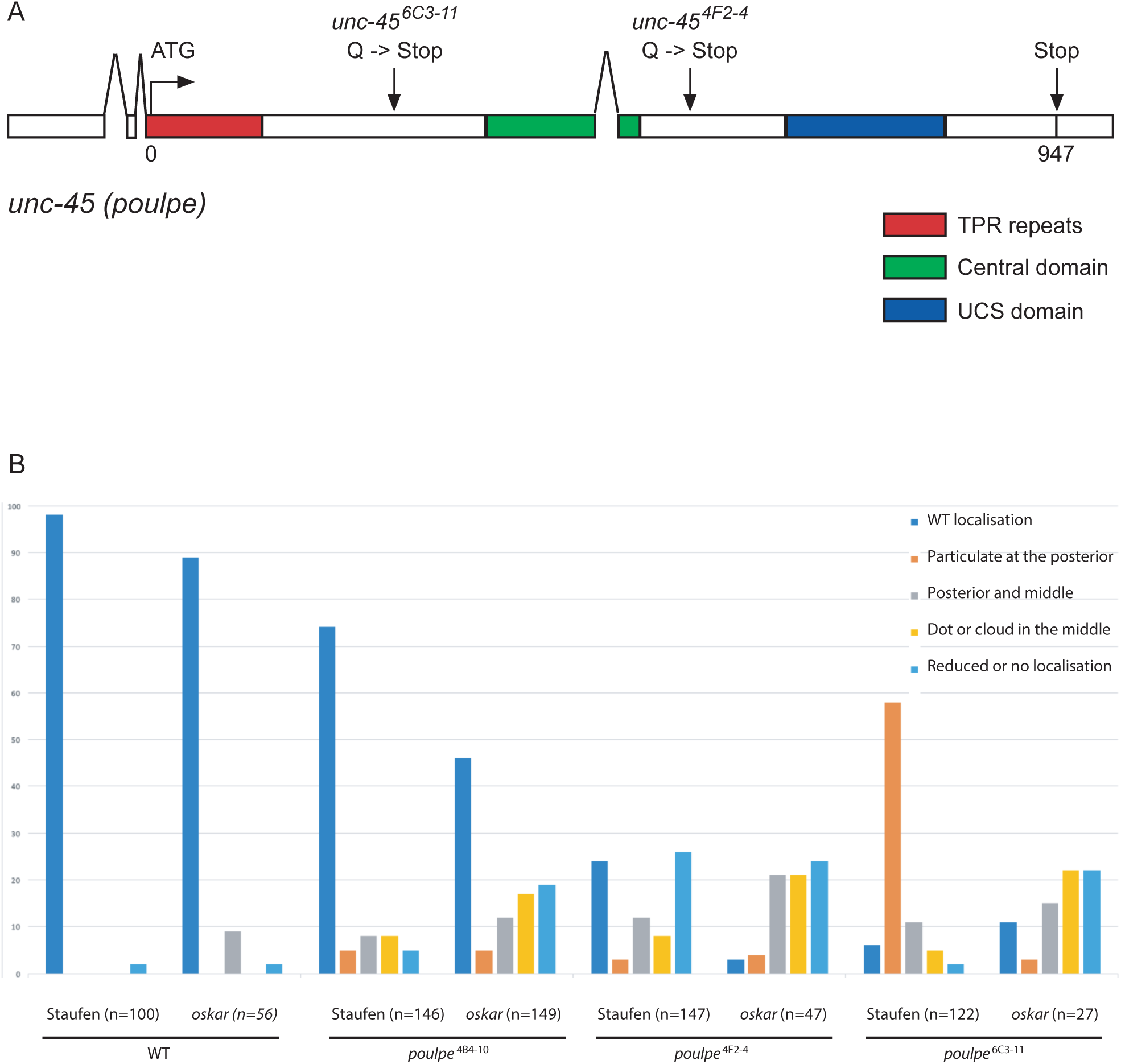
Localisation of the unc-45 mutations and their phenotypes in the oocyte. (A) Schematic representation of the *unc-45* (*poulpe*) coding region showing the positions of the nonsense mutations in *poulpe [6C31-11]* and *poulpe [4F2-4]*. TPR: tetratricopeptide repeat (red), Central domain (green) and UCS: UNC-45, CRO1, She4p (blue) (B) Quantification of the Staufen and *oskar* mRNA localisation defects in WT oocytes andoocytes mutant or the *poulpe [4B4-10], [4F2-4]* and *[6C3-11]* . Normal localisation to the posterior cortex of the oocyte (dark blue), Particulate at the oocyte posterior (orange), Posterior and middle of the oocyte(grey), Dot or cloud in the middle of teh oocyte (yellow), Reduced or no localisation (light blue).

**Supplementary Figure 2.**
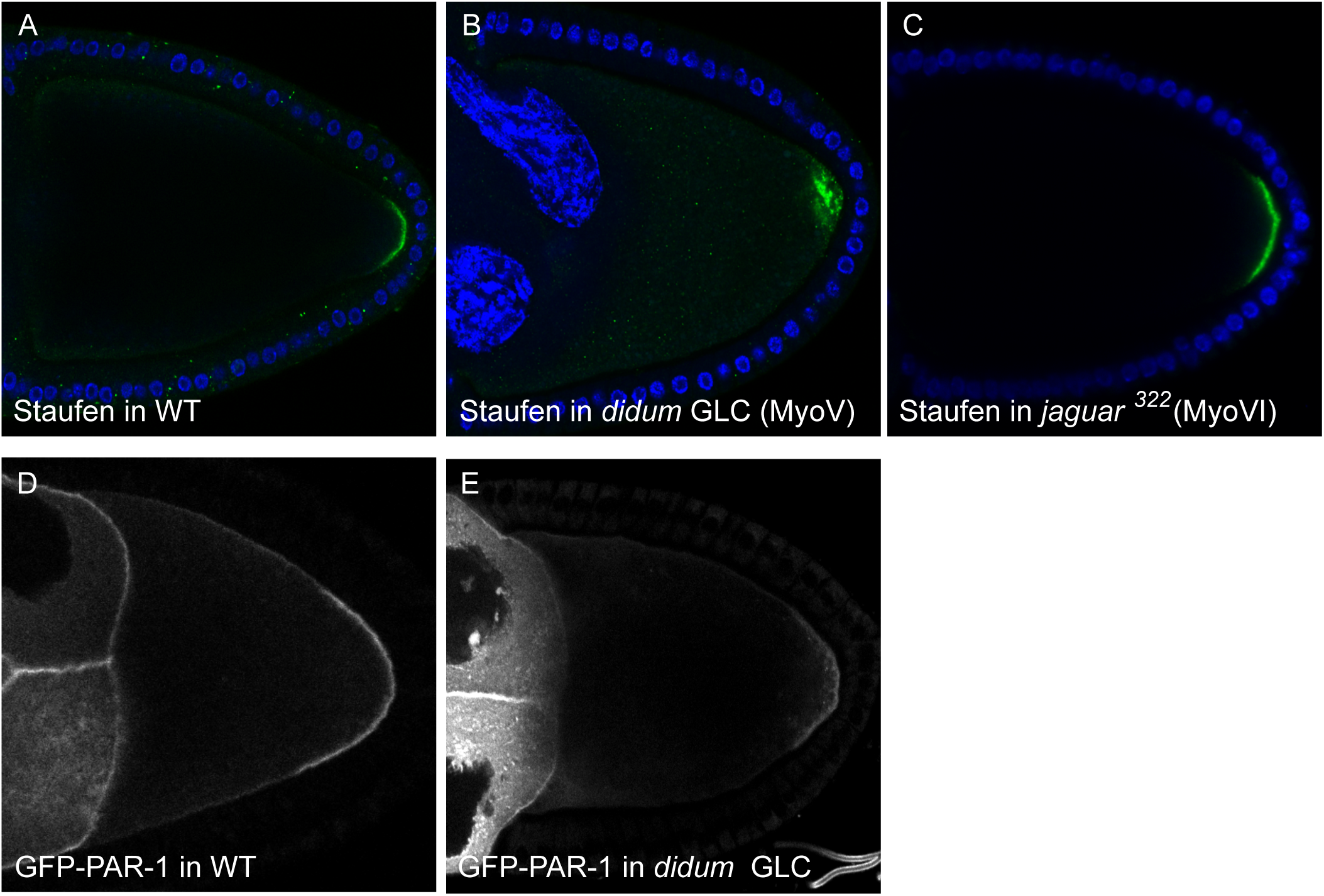
MyosinV acts downstrem of Par-1. (A) A confocal image of a wild-type egg chamber showing the localisation of Staufen (green) in a crescent at the posterior cortex of the oocyte; DAPI (blue). (B) A confocal image showing Staufen localisation in a *didum* (Myosin V) mutant germline clone (GLC). Staufen forms a diffuse cloud near the posterior pole; Staufen (green) and DAPI (blue). (C) A confocal image showing Staufen localisation in a *jaguar^322^* (Myosin VI) homozygous mutant. Staufen forms a normal crescent at the posterior pole of the oocyte; Staufen (green) and DAPI (blue). (D-E) Confocal images showing Par-1 GFP in a crescent at the posterior cortex of the oocyte both in a wild type oocyte (D) and in *didum* (Myosin V) homozygous mutant oocyte (E).

**Supplementary figure 3.**
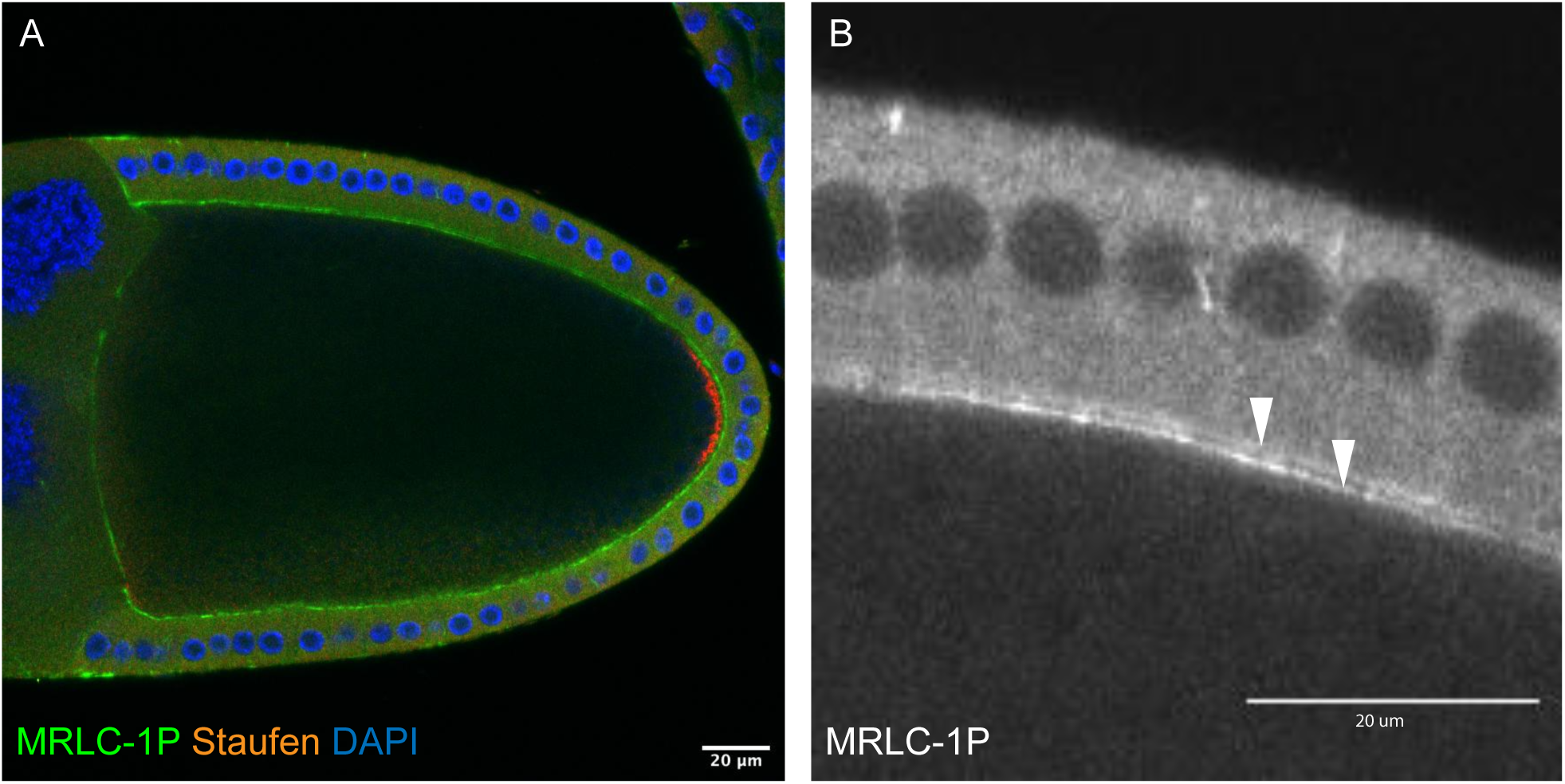
Mono-phosphorylated MyoII localised uniformly along the oocyte cortex. (A) Antibody staining of mono-phosphorylated MyoII (MRLC-1P) (green) and Staufen (red) in a wild-type oocyte; DAPI (blue). MRLC-1P is distributed uniformly along the oocyte cortex. (A’) Close up of the mono-phosphorylated MyoII (MRLC-1P) showing that MRLC-1P is localised at the apical side of the follicle cells and along the oocyte cortex (white arrows).

**Supplementary Figure 4.**
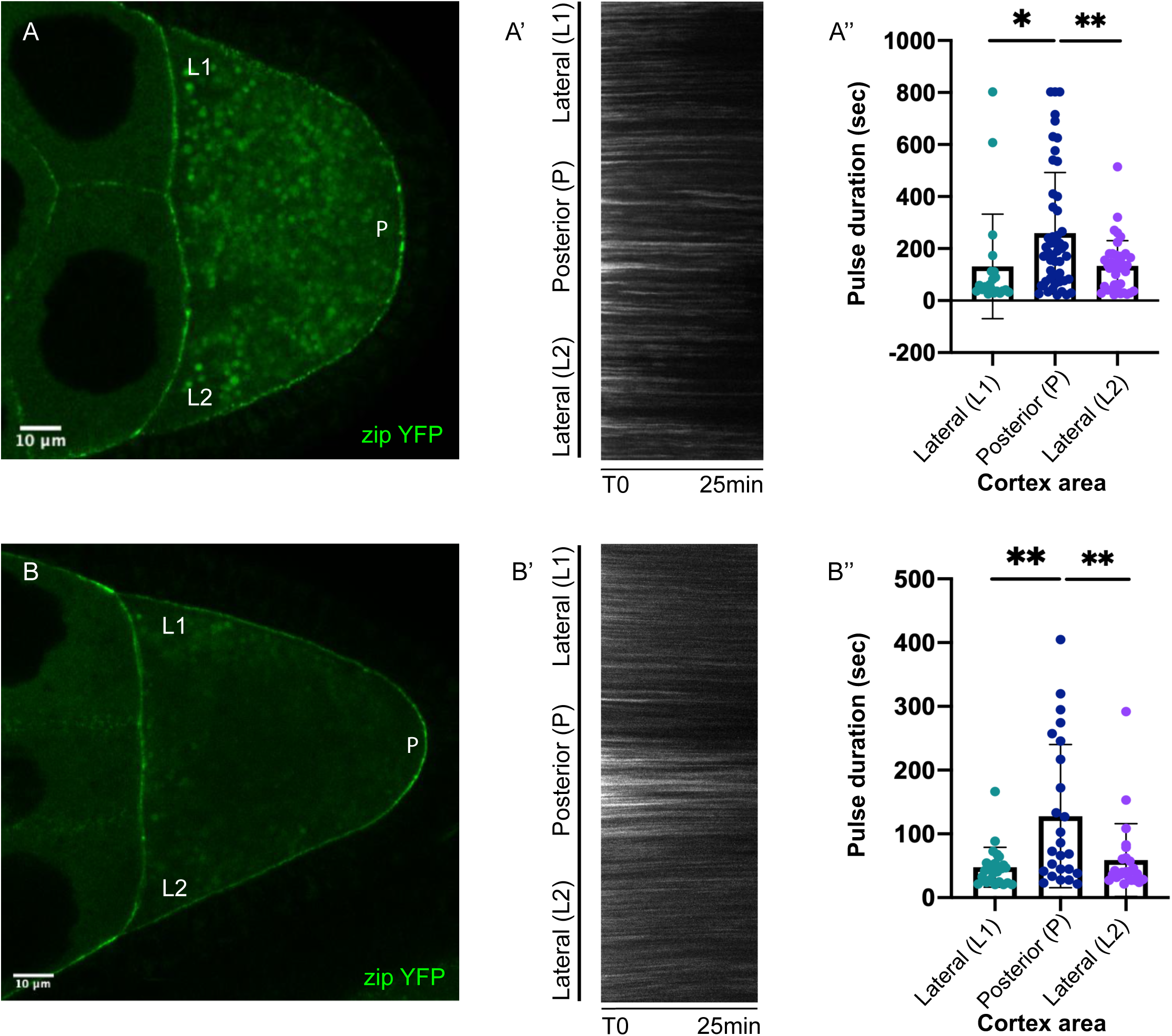
MyoII forms cortical foci that persist longer at the posterior – Additional examples. (A-B) Still images from movies of Zipper-GFP in stage 9 oocytes. (A’-B’) Kymographs showing the changes to Zipper-GFP levels over time along the cortex. Zipper-GFP foci remain stationary and oscillate in intensity over time. (A’’-B’’) A graph showing the durations of Zipper-GFP pulses at the lateral and posterior cortex (L1, P, L2). Pulse durations are measured using an automated detection and segmentation algorithm. The pulses at the posterior last twice as long as the lateral pulses.

## References

[1] Goldstein B, Hird SN. Specification of the anteroposterior axis in *Caenorhabditis elegans*. Development 1996;122:1467–74.

[2] Kapoor S, Kotak S. Centrosome Aurora A regulates RhoGEF ECT-2 localisation and ensures a single PAR-2 polarity axis in *C. elegans* embryos. Development 2019;146:dev174565. doi:10.1242/dev.174565.

[3] Klinkert K, Levernier N, Gross P, Gentili C, Tobel von L, Pierron M, et al. Aurora A depletion reveals centrosome-independent polarization mechanism in *Caenorhabditis elegans*. Elife 2019;8:316. doi:10.7554/eLife.44552.

[4] Zhao P, Teng X, Tantirimudalige SN, Nishikawa M, Wohland T, Toyama Y, et al. Aurora-A breaks symmetry in contractile actomyosin networks independently of its role in centrosome maturation. Dev Cell 2019;48:631–6. doi:10.1016/j.devcel.2019.02.012.

[5] Sailer A, Anneken A, Li Y, Lee S, Munro E. Dynamic opposition of clustered proteins stabilizes cortical polarity in the *C. elegans* zygote. Dev Cell 2015;35:131–42. doi:10.1016/j.devcel.2015.09.006.

[6] Dickinson DJ, Schwager F, Pintard L, Gotta M, Goldstein B. A single-cell biochemistry approach reveals PAR complex dynamics during cell polarization. Dev Cell 2017;42:416–434.e11. doi:10.1016/j.devcel.2017.07.024.

[7] Rodriguez J, Peglion F, Martin J, Hubatsch L, Reich J, Hirani N, et al. aPKC cycles between functionally distinct PAR protein assemblies to drive cell polarity. Dev Cell 2017;42:400–9. doi:10.1016/j.devcel.2017.07.007.

[8] Lang CF, Munro E. The PAR proteins: from molecular circuits to dynamic self-stabilizing cell polarity. Development 2017;144:3405–16. doi:10.1242/dev.139063.

[9] Bastock R, St Johnston D. *Drosophila* oogenesis. Curr Biol 2008;18:R1082–7. doi:10.1016/j.cub.2008.09.011.

[10] Godt D, Tepass U. *Drosophila* oocyte localisation is mediated by differential cadherin-based adhesion. Nature 1998;395:387–91.

[11] González-Reyes A, St Johnston D. The *Drosophila* AP axis is polarised by the cadherin-mediated positioning of the oocyte. Development 1998;125:3635–44.

[12] Torres I, Lopez-Schier H, St Johnston D. A Notch/Delta-dependent relay mechanism establishes anterior-posterior polarity in *Drosophila*. Dev Cell 2003;5:547–58.

[13] González-Reyes A, St Johnston D. Patterning of the follicle cell epithelium along the anterior-posterior axis during *Drosophila* oogenesis. Development 1998;125:2837– 46.

[14] Xi R, McGregor JR, Harrison DA. A gradient of JAK pathway activity patterns the anterior-posterior axis of the follicular epithelium. Dev Cell 2003;4:167–77.

[15] González-Reyes A, Elliott H, St Johnston D. Polarization of both major body axes in *Drosophila* by *gurken*-*torpedo* signalling. Nature 1995;375:654–8. doi:10.1038/375654a0.

[16] Roth S, Neuman-Silberberg FS, Barcelo G, Schüpbach T. Cornichon and the EGF receptor signaling process are necessary for both anterior-posterior and dorsal-ventral pattern formation in *Drosophila*. Cell 1995;81:967–78. doi:10.1016/0092-8674(95)90016-0.

[17] Wittes J, Schupbach T. A gene expression screen in *Drosophila melanogaster* identifies novel JAK/STAT and EGFR targets during oogenesis. G3 (Bethesda) 2019;9:47–60. doi:10.1534/g3.118.200786.

[18] Shulman JM, Benton R, St Johnston D. The Drosophila homolog of *C. elegans* PAR-1 organizes the oocyte cytoskeleton and directs *oskar* mRNA localization to the posterior pole. Cell 2000;101:377–88. doi:10.1016/s0092-8674(00)80848-x.

[19] Tomancak P, Piano F, Riechmann V, Gunsalus KC, Kemphues KJ, Ephrussi A. A *Drosophila melanogaster* homologue of *Caenorhabditis elegans par-1* acts at an early step in embryonic-axis formation. Nature Cell Biol 2000;2:458–60. doi:10.1038/35017101.

[20] Doerflinger H, Benton R, Torres IL, Zwart MF, St Johnston D. *Drosophila* anterior-posterior polarity requires actin-dependent PAR-1 recruitment to the oocyte posterior. Curr Biol 2006;16:1090–5. doi:10.1016/j.cub.2006.04.001.

[21] Doerflinger H, Vogt N, Torres IL, Mirouse V, Koch I, Nüsslein-Volhard C, et al. Bazooka is required for polarisation of the *Drosophila* anterior-posterior axis. Development 2010;137:1765–73. doi:10.1242/dev.045807.

[22] Benton R, St Johnston D. *Drosophila* PAR-1 and 14-3-3 inhibit Bazooka/PAR-3 to establish complementary cortical domains in polarized cells. Cell 2003;115:691–704.

[23] Zimyanin VL, Belaya K, Pecreaux J, Gilchrist MJ, Clark A, Davis I, et al. In vivo imaging of *oskar* mRNA transport reveals the mechanism of posterior localization. Cell 2008;134:843–53. doi:10.1016/j.cell.2008.06.053.

[24] Nashchekin D, Fernandes AR, St Johnston D. Patronin/Shot cortical foci assemble the noncentrosomal microtubule array that specifies the *Drosophila* anterior-posterior Axis. Dev Cell 2016;38:61–72. doi:10.1016/j.devcel.2016.06.010.

[25] Morais-de-Sá E, Mukherjee A, Lowe N, St Johnston D. Slmb antagonises the aPKC/Par-6 complex to control oocyte and epithelial polarity. Development 2014;141:2984–92. doi:10.1242/dev.109827.

[26] Martin SG, Leclerc V, Smith-Litière K, St Johnston D. The identification of novel genes required for *Drosophila* anteroposterior axis formation in a germline clone screen using GFP-Staufen. Development 2003;130:4201–15.

[27] Clark I, Giniger E, Ruohola-Baker H, L Y Jan, Y N Jan. Transient posterior localization of a kinesin fusion protein reflects anteroposterior polarity of the *Drosophila* oocyte. Curr Biol 1994;4:289–300.

[28] Parton RM, Hamilton RS, Ball G, Yang L, Cullen CF, Lu W, et al. A PAR-1-dependent orientation gradient of dynamic microtubules directs posterior cargo transport in the Drosophila oocyte. J Cell Biol 2011;194:121–35. doi:10.1083/jcb.201103160.

[29] Bellen HJ, Levis RW, Liao G, He Y, Carlson JW, Tsang G, et al. The BDGP gene disruption project: single transposon insertions associated with 40% of *Drosophila* genes. Genetics 2004;167:761–81. doi:10.1534/genetics.104.026427.

[30] Melkani GC, Bodmer R, Ocorr K, Bernstein SI. The UNC-45 chaperone is critical for establishing myosin-based myofibrillar organization and cardiac contractility in the *Drosophila* heart model. PLoS ONE 2011;6:e22579. doi:10.1371/journal.pone.0022579.

[31] Lee CF, Melkani GC, Bernstein SI. The UNC-45 myosin chaperone: from worms to flies to vertebrates. Int Rev Cell Mol Biol 2014;313:103–44. doi:10.1016/B978-0-12-800177-6.00004-9.

[32] Krauss J, López de Quinto S, Nüsslein-Volhard C, Ephrussi A. Myosin-V regulates *oskar* mRNA localization in the *Drosophila* oocyte. Curr Biol 2009;19:1058–63. doi:10.1016/j.cub.2009.04.062.

[33] Lu W, Lakonishok M, Liu R, Billington N, Rich A, Glotzer M, et al. Competition between kinesin-1 and myosin-V defines *Drosophila* posterior determination. Elife 2020;9:779. doi:10.7554/eLife.54216.

[34] Okumura T, Sasamura T, Inatomi M, Hozumi S, Nakamura M, Hatori R, et al. Class I myosins have overlapping and specialized functions in left-right asymmetric development in Drosophila. Genetics 2015;199:1183–99. doi:10.1534/genetics.115.174698.

[35] Cheeks RJ, Canman JC, Gabriel WN, Meyer N, Strome S, Goldstein B. *C. elegans* PAR proteins function by mobilizing and stabilizing asymmetrically localized protein complexes. Curr Biol 2004;14:851–62. doi:10.1016/j.cub.2004.05.022.

[36] Munro E, Nance J, Priess JR. Cortical flows powered by asymmetrical contraction transport PAR proteins to establish and maintain anterior-posterior polarity in the early *C. elegans* embryo. Dev Cell 2004;7:413–24. doi:10.1016/j.devcel.2004.08.001.

[37] Vicente-Manzanares M, Ma X, Adelstein RS, Horwitz AR. Non-muscle myosin II takes centre stage in cell adhesion and migration. Nat Rev Mol Cell Biol 2009;10:778–90. doi:10.1038/nrm2786.

[38] Lowe N, Rees JS, Roote J, Ryder E, Armean IM, Johnson G, et al. Analysis of the expression patterns, subcellular localisations and interaction partners of *Drosophila* proteins using a pigP protein trap library. Development 2014;141:3994–4005. doi:10.1242/dev.111054.

[39] Jordan P, Karess R. Myosin light chain-activating phosphorylation sites are required for oogenesis in *Drosophila*. J Cell Biol 1997;139:1805–19.

[40] Heissler SM, Sellers JR. Myosin light chains: Teaching old dogs new tricks. Bioarchitecture 2014;4:169–88. doi:10.1080/19490992.2015.1054092.

[41] Vasquez CG, Heissler SM, Billington N, Sellers JR, Martin AC. *Drosophila* non-muscle myosin II motor activity determines the rate of tissue folding. Elife 2016;5:1761. doi:10.7554/eLife.20828.

[42] Zhang L, Ward RE. Distinct tissue distributions and subcellular localizations of differently phosphorylated forms of the myosin regulatory light chain in *Drosophila*. Gene Expr Patterns 2011;11:93–104. doi:10.1016/j.gep.2010.09.008.

[43] Vasquez CG, Tworoger M, Martin AC. Dynamic myosin phosphorylation regulates contractile pulses and tissue integrity during epithelial morphogenesis. J Cell Biol 2014;206:435–50. doi:10.1083/jcb.201402004.

[44] Dawes-Hoang RE, Parmar KM, Christiansen AE, Phelps CB, Brand AH, Wieschaus EF. Folded gastrulation, cell shape change and the control of myosin localization. Development 2005;132:4165–78. doi:10.1242/dev.01938.

[45] Goehring NW, Trong PK, Bois JS, Chowdhury D, Nicola EM, Hyman AA, et al. Polarization of PAR Proteins by advective triggering of a pattern-forming system. Science 2011;334:1137–41. doi:10.1126/science.1208619.

[46] Martin AC, Kaschube M, Wieschaus EF. Pulsed contractions of an actin-myosin network drive apical constriction. Nature 2009;457:495–9. doi:10.1038/nature07522.

[47] He L, Wang X, Tang HL, Montell DJ. Tissue elongation requires oscillating contractions of a basal actomyosin network. Nat Cell Biol 2010;12:1133–42. doi:10.1038/ncb2124.

[48] Mason FM, Tworoger M, Martin AC. Apical domain polarization localizes actin-myosin activity to drive ratchet-like apical constriction. Nat Cell Biol 2013;15:926– 36. doi:10.1038/ncb2796.

[49] Alégot H, Pouchin P, Bardot O, Mirouse V. Jak-Stat pathway induces *Drosophila* follicle elongation by a gradient of apical contractility. Elife 2018;7:e32943. doi:10.7554/eLife.32943.

[50] Maekawa M, Ishizaki T, Boku S, Watanabe N, Fujita A, Iwamatsu A, et al. Signaling from Rho to the actin cytoskeleton through protein kinases ROCK and LIM-kinase. Science 1999;285:895–8. doi:10.1126/science.285.5429.895.

[51] Kaibuchi K, Kuroda S, Amano M. Regulation of the cytoskeleton and cell adhesion by the Rho family GTPases in mammalian cells. Annual Review of Biochemistry 1999;68:459–86. doi:10.1146/annurev.biochem.68.1.459.

[52] Munjal A, Philippe J-M, Munro E, Lecuit T. A self-organized biomechanical network drives shape changes during tissue morphogenesis. Nature 2015;524:351–5. doi:10.1038/nature14603.

[53] del Rio A, Perez-Jimenez R, Liu R, Roca-Cusachs P, Fernandez JM, Sheetz MP. Stretching single talin rod molecules activates vinculin binding. Science 2009;323:638–41. doi:10.1126/science.1162912.

[54] Yonemura S, Wada Y, Watanabe T, Nagafuchi A, Shibata M. alpha-Catenin as a tension transducer that induces adherens junction development. Nat Cell Biol 2010;12:533–42. doi:10.1038/ncb2055.

[55] Li T, Giagtzoglou N, Eberl DF, Jaiswal SN, Cai T, Godt D, et al. The E3 ligase Ubr3 regulates Usher syndrome and MYH9 disorder proteins in the auditory organs of *Drosophila* and mammals. Elife 2016;5:347. doi:10.7554/eLife.15258.

[56] Guo S, Kemphues KJ. A non-muscle myosin required for embryonic polarity in *Caenorhabditis elegans*. Nature 1996;382:455–8. doi:10.1038/382455a0.

[57] Rolls MM, Albertson R, Shih H-P, Lee C-Y, Doe CQ. *Drosophila* aPKC regulates cell polarity and cell proliferation in neuroblasts and epithelia. J Cell Biol 2003;163:1089–98. doi:10.1083/jcb.200306079.

[58] Atwood SX, Prehoda KE. aPKC phosphorylates Miranda to polarize fate determinants during neuroblast asymmetric cell division. Curr Biol 2009;19:723–9. doi:10.1016/j.cub.2009.03.056.

[59] Hannaford MR, Ramat A, Loyer N, Januschke J. aPKC-mediated displacement and actomyosin-mediated retention polarize Miranda in *Drosophila* neuroblasts. Elife 2018;7:166. doi:10.7554/eLife.29939.

[60] Hannaford M, Loyer N, Tonelli F, Zoltner M, Januschke J. A chemical-genetics approach to study the role of atypical Protein Kinase C in *Drosophila*. Development 2019;146:dev170589. doi:10.1242/dev.170589.

[61] Schnorrer F, Luschnig S, Koch I, Nüsslein-Volhard C. Gamma-tubulin37C and gamma-tubulin ring complex protein 75 are essential for *bicoid* RNA localization during *Drosophila* oogenesis. Dev Cell 2002;3:685–96.

[62] Perkins LA, Holderbaum L, Tao R, Hu Y, Sopko R, McCall K, et al. The Transgenic RNAi Project at Harvard Medical School: Resources and Validation. Genetics 2015;201:843–52. doi:10.1534/genetics.115.180208.

[63] Gloor G, Nassif N, Johnson-Schlitz D, Preston C, Engels W. Targeted gene replacement in *Drosophila* via P element-induced gap repair. Science 1991;253:1110–7. doi:10.1126/science.1653452.

